# Rapid suppression and sustained activation of distinct cortical regions for a delayed sensory-triggered motor response

**DOI:** 10.1101/2020.10.06.326678

**Authors:** Vahid Esmaeili, Keita Tamura, Samuel P. Muscinelli, Alireza Modirshanechi, Marta Boscaglia, Ashley B. Lee, Anastasiia Oryshchuk, Georgios Foustoukos, Yanqi Liu, Sylvain Crochet, Wulfram Gerstner, Carl C.H. Petersen

**Author notes:** Contributed equally to this study (Vahid Esmaeili, Keita Tamura). Lead contact: Carl Petersen, Laboratory of Sensory Processing, Brain Mind Institute, Faculty of Life Sciences, SV-BMI-LSENS Station 19, Ecole Polytechnique Fédérale de Lausanne (EPFL), CH-1015 Lausanne, Switzerland.

## Abstract

The neuronal mechanisms generating a delayed motor response initiated by a sensory cue remain elusive. Here, we tracked the precise sequence of cortical activity in mice transforming a brief whisker stimulus into delayed licking using wide-field calcium imaging, multi-region high-density electrophysiology and time-resolved optogenetic manipulation. Rapid activity evoked by whisker deflection acquired two prominent features for task performance: i) an enhanced excitation of secondary whisker motor cortex, suggesting its important role connecting whisker sensory processing to lick motor planning, and ii) a transient reduction of activity in orofacial sensorimotor cortex, which contributed to suppressing premature licking. Subsequent widespread cortical activity during the delay period largely correlated with anticipatory movements, but when these were accounted for, a focal sustained activity remained in frontal cortex, which was causally essential for licking in the response period. Our results demonstrate key cortical nodes for motor plan generation and timely execution in delayed goal-directed licking.

## INTRODUCTION

Incoming sensory information is processed in a learning- and context-dependent manner to direct behavior. Timely execution of appropriate action requires motor planning, in particular when the movement triggered by a sensory cue needs to be delayed. In this situation, the motor plan must persist throughout the delay period while the immediate execution of the motor response needs to be suppressed. Delayed-response paradigms are often used to study the neuronal circuits of sensorimotor transformation because they allow to temporally isolate the neuronal activity that bridges sensation and action. In such paradigms, prominent delay period activity has been reported in many cortical areas (Chabrol et al., 2019; Chen et al., 2017; Erlich et al., 2011; Esmaeili and Diamond, 2019; Fassihi et al., 2017; Funahashi et al., 1989; Fuster and Alexander, 1971; Gilad et al., 2018; Guo et al., 2014; Li et al., 2015; Makino et al., 2017; Tanji and Evarts, 1976). In particular, a previous study in mice identified delay period activity in the anterolateral motor (ALM) cortex, which causally contributed to a lick motor plan (Guo et al., 2014). The persistent delay period activity in ALM is driven through a recurrent thalamocortical loop (Guo et al., 2017), and further supported by cerebellar interactions (Chabrol et al., 2019; Gao et al., 2018). The circuit mechanisms maintaining the persistent activity in ALM are therefore beginning to be understood. However, less is known about the circuits that initiate such persistent activity, and how task learning shapes such circuits. In addition, how the persistent neuronal activity is related to body movements which animals often exhibit during delay periods needs to be carefully considered (Musall et al., 2019; Steinmetz et al., 2019; Stringer et al., 2019). Similarly, the neuronal circuits contributing to withholding a premature motor response during the delay are poorly understood. To dissect this process, it would be crucial to examine how neuronal activity flows across brain areas as sensory information is transformed into goal-directed motor plans (de Lafuente and Romo, 2006), and to investigate how the underlying sensory and motor circuits become connected through reward-based learning (Esmaeili et al., 2020).

Here, we address these questions in head-restrained mice performing a whisker-detection task with delayed licking to report perceived stimuli. In our task, a brief and well-defined sensory input is rapidly transformed into a decision and mice need to withhold the response until the end of the delay period. Through a unified and comprehensive approach, we detail the spatiotemporal map of causal cortical processing which emerges across learning. We found that following the fast sensory-evoked response in somatosensory cortex (Petersen, 2019), the activity in orofacial sensorimotor cortex (Mayrhofer et al., 2019) was rapidly and transiently suppressed, which contributed causally to withholding premature licking. The subsequent rapid sequential excitation of frontal cortical regions and their changes across task learning revealed that secondary whisker motor cortex (wM2) likely plays a key role linking whisker sensation to motor planning. We also found that the global activation of dorsal cortex during the delay period could be largely ascribed to preparatory movements that develop with learning, except for a localized neuronal activity in ALM (Komiyama et al., 2010), consistent with previous studies (Chen et al., 2017; Guo et al., 2014). Our results therefore point to task epoch-specific contributions of distinct cortical regions to whisker-triggered planning of goal-directed licking and timely execution of planned lick responses.

## RESULTS

### Behavioral changes accompanying delayed-response task learning

We designed a Go/No-Go learning paradigm where head-restrained mice learned to lick in response to a whisker stimulus after a one second delay period (Figure 1A-C). To precisely track the sequence of cortical responses, we used a single, short (10 ms) deflection of the right C2 whisker. Trial start was indicated by a 200 ms light flash, followed 1 s later by the whisker stimulus in 50% of the trials (referred to as Go trials); after a subsequent 1 s delay, a 200 ms auditory tone signaled the beginning of a 1 s response window. Licking during the response window, in Go trials, was rewarded with a drop of water, whereas licking before the auditory tone (early lick) led to abortion of the trial and time-out punishment (Figure 1B). To study essential neuronal circuit changes specific to the coupling of the whisker stimulus with the licking response, a two-phase learning paradigm was implemented: i) Pretraining, and ii) Whisker training (Figure 1C). Pretraining included trials with visual and auditory cues only, and licking during the response window was rewarded, while licking before the auditory cue aborted the trial. Novice mice only went through the Pretraining, which established the general task structure. Expert mice followed an additional Whisker training phase, during which they learned the final task structure (Figures 1B-C and S1A).

**Figure 1.**
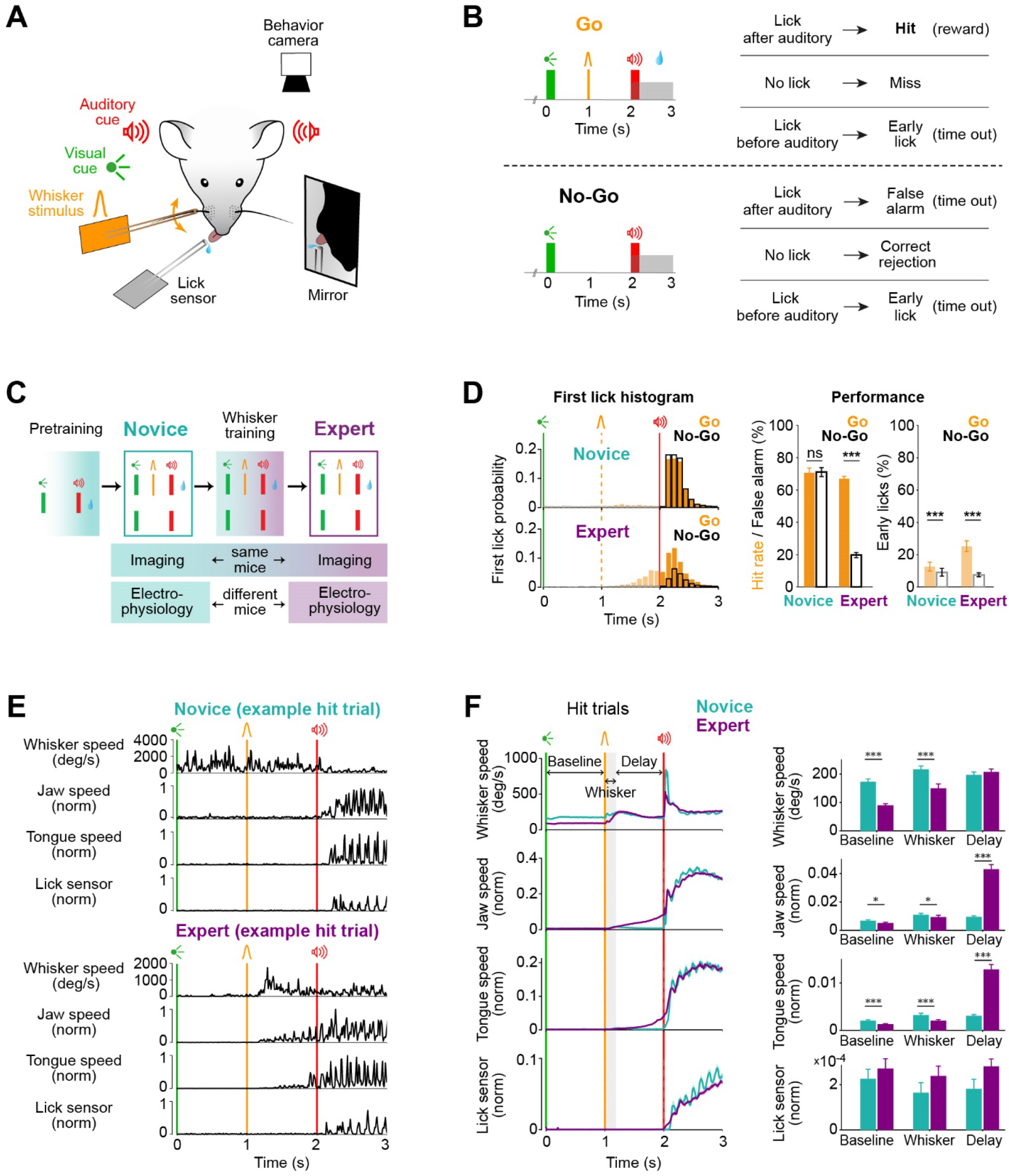
Learning a whisker detection task with a delayed response changes licking patterns and orofacial movements. (A and B) Delayed whisker detection task. A, Behavioral setup. Sensory stimuli were delivered to head-restrained mice and licking and orofacial movements were monitored using a piezoelectric lick sensor and a behavior camera. B, Task structure and trial outcomes in Go and No-Go trials. (C) Learning paradigm. All mice went through visual-auditory Pretraining, where all licks after the auditory cue were rewarded. Expert mice went through Whisker training where final task structure was used as in panel B. Neural data were obtained using the final task design in Novice mice and Expert mice, before and after Whisker training respectively. The same mice were imaged during Novice and Expert stages, while different mice were used for electrophysiological recordings. (D) Task performance. *Left*, first lick time histogram was similar in Go vs No-Go trials in Novice mice, but differed in Expert mice. Early licks (licks between visual cue and auditory cue) are shown with light colors. *Middle*, Novice mice licked equally in Go and No-Go trials, whereas Expert mice licked preferentially in Go trials (quantified as mean ± SEM across all completed trials; Novice, n=15 mice; Expert, n=25 mice). *Right*, both groups of mice made more early licks in Go compared to No-Go trials. *** indicates *p*<0.001 according to Wilcoxon signed-rank test. (E and F) Orofacial movements. E, Example trials from Novice (*top*) and Expert (*bottom*) mice: extracted angular speed of left C2-whisker, and normalized jaw and tongue speed together with the lick sensor signal are plotted along the trial time-course. F, Average movement (mean ± SEM) traces (*left*) and bar plots comparing Novice and Expert movements in 3 different windows (*right*). Expert mice reduced all movements during Baseline and Whisker time windows, while they increased tongue and jaw movements during the Delay. Asterisks represent statistical comparison of movement between Novice and Expert mice in: Baseline (0-1 s), Whisker (1-1.2 s), and Delay (1.2-2 s) time windows (Wilcoxon rank-sum test, false discovery rate (FDR) corrected, ***: *p*<0.001, *: *p*<0.05). Only mice used for electrophysiology (8 Novice and 18 Expert mice) are plotted. See also Figure S1.

Novice and Expert mice were recorded in the same final task condition, but performed differently. While, Novice mice licked in both Go and No-Go trials, Expert mice had learned to lick preferentially in Go trials (Figures 1D and S1A; mean ± SEM, Novice: Hit=70.6 ± 3%, False-alarm=71.1 ± 2.7%, *p*=0.85, n=15 mice; Expert: Hit=67± 1.5%, False-alarm=19.7 ± 1.6%, *p*<0.001, n=25 mice; Wilcoxon signed-rank test). Expert mice made more frequent premature early licks in Go-trials compared to Novice mice (Figures 1D; mean ± SEM, Novice=12.5 ± 2.7%, Expert= 25.1 ± 3.3%, *p*=0.02; Wilcoxon rank-sum test) and most of their early licks happened toward the end of the delay period, reflecting predictive licking. Considering trials with licking during the response window, Expert mice showed longer reaction times in No-Go trials (False-alarm) compared to Go trials (Hit) (Figure S1B; mean ± SEM, Novice: Hit=298.5 ± 21.2 ms, False-alarm=292.7 ± 21.4 ms, *p*=0.14; Expert: Hit=297.8 ± 16.7 ms, False-alarm=380.3 ± 16.3 ms, *p*<0.01; Wilcoxon signed-rank test). These results indicate that Expert mice used whisker information and learned to produce delayed licking. After Whisker training, mice also adopted new movement strategies (Figures 1E-F and S1C-D). In Hit trials, Expert mice compared to Novice mice decreased whisker movement before whisker stimulus, possibly to improve the detection of brief whisker stimuli in the receptive mode of perception (Diamond and Arabzadeh, 2013; Kyriakatos et al., 2017). The tongue and jaw movements in the delay period after the whisker stimulus increased in Hit trials of Expert mice compared to Novice mice, reflecting preparation for licking. These anticipatory movements were absent in Miss and Correct-rejection trials (Figure S1C), and thus correlated with the perceptual response. These patterns were similar comparing mice used for electrophysiology and imaging (Figures 1F and S1D).

### Emergence of cortical activation and deactivation patterns through whisker training

The delay-task enables the investigation of different aspects of neuronal computations underlying reward-based behavior including: sensory processing, motor planning and motor execution in well-isolated time windows. As a first step, we mapped the large-scale dynamics of cortical activity using wide-field calcium imaging at a high temporal resolution (100 frames per second) (Figures 2, S2 and S3). In transgenic mice expressing a fluorescent calcium indicator in pyramidal neurons (RCaMP mice) (Bethge et al., 2017), functional images of the left dorsal cortex were obtained through an intact skull preparation, and registered to the Allen Mouse Brain Common Coordinate Framework (Figures 2A-B) (Lein et al., 2007; Wang et al., 2020).

**Figure 2.**
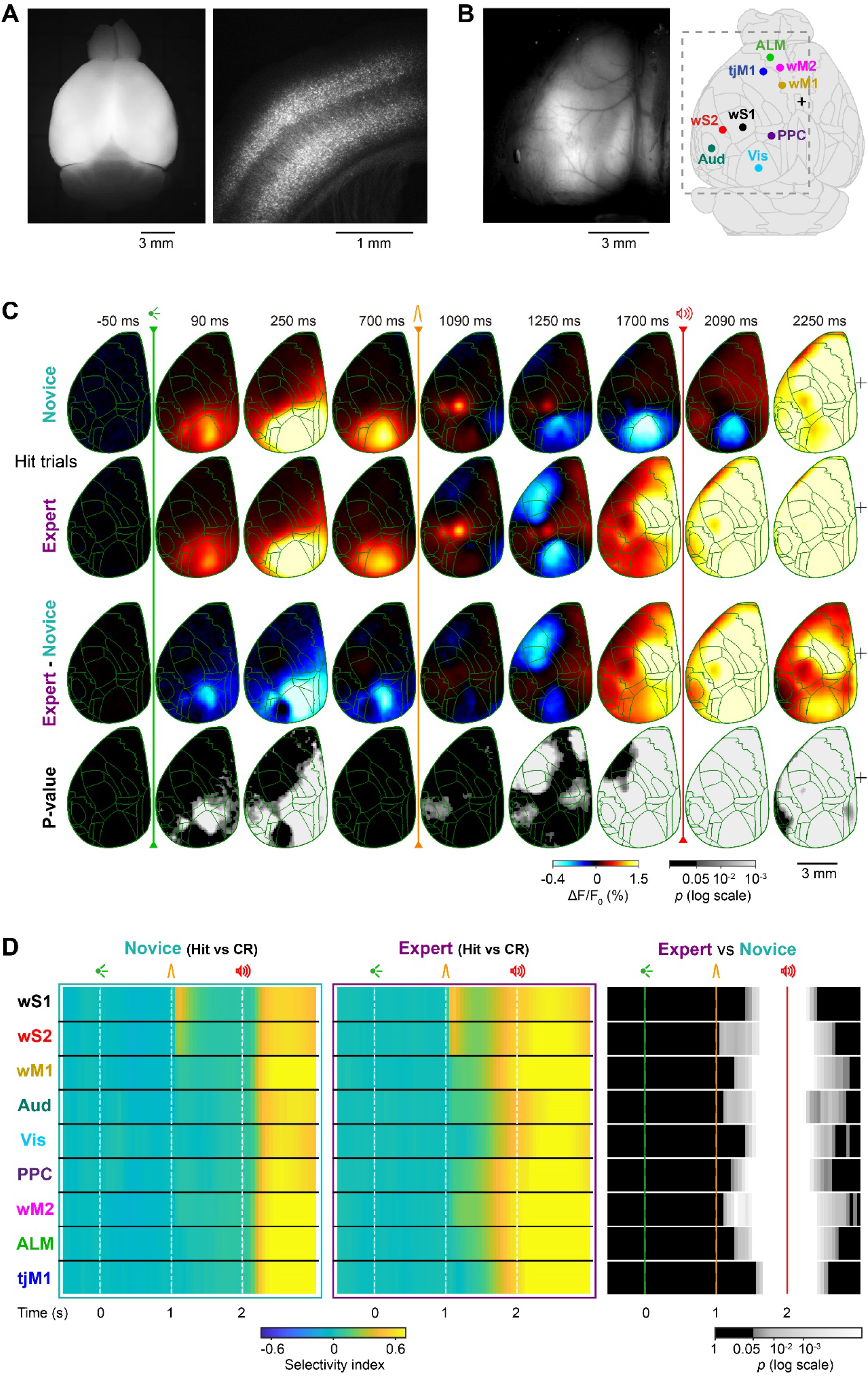
Wide-field imaging reveals global changes in cortical processing. (A and B) Wide-field calcium imaging in Emx1-RCaMP mice. A, Fluorescence images of an ex vivo fixed brain in dorsal view (*left*) and a coronal section of somatosensory area (*right*) showing RCaMP expression in pyramidal neurons in deep and superficial layers. B, In vivo fluorescence image of the tilted (24 degrees) left dorsal hemisphere through a transparent, intact skull preparation (*left*). In vivo images were registered to the Allen Mouse Brain Atlas (*right*). Cortical areas targeted for electrophysiological experiments are indicated: wS1, primary whisker somatosensory cortex; wS2, secondary whisker somatosensory cortex; wM1, primary whisker motor cortex; wM2, secondary whisker motor cortex; Aud, auditory area; Vis, visual area; PPC, posterior parietal cortex; tjM1, primary tongue and jaw motor cortex; ALM, anterior lateral motor area. (C) Grand-average time-course of global cortical activity in Hit trials for Novice vs Expert mice. Each frame shows ΔF/F_0_ without temporal smoothing (10 ms/frame). For each pixel, baseline activity in a 50 ms window before visual cue onset was subtracted. Mean calcium activity of 62 Novice and 82 Expert sessions from 7 mice, Novice and Expert difference, and the statistical significance of the difference (*p*-value of Wilcoxon rank-sum test, FDR-corrected, *p*<0.05) are plotted from top to bottom. Green traces, anatomical borders based on Allen Mouse Brain Atlas. Black ‘+’ indicates bregma. (D) Selectivity index in Novice and Expert mice. For each brain region, selectivity between Hit versus Correct-rejection trials was determined in non-overlapping 50 ms bins based on the area under the ROC curve. Mean selectivity of each area in 62 Novice and 82 Expert sessions from 7 mice, and the statistical significance of the difference (*p*-value of non-parametric permutation test, FDR-corrected, *p*<0.05) is plotted. ROI size, 3×3 pixels. See also Figures S2-S3 and Videos S1-S4.

To examine the changes in cortical processing upon learning, we compared the activity in the same mice (n=7) before (Novice, 62 sessions) and after (Expert, 82 sessions) whisker training (Figures 2C for Hit trials and S2A for Correct-rejection trials, Videos S1-2). The visual cue evoked responses in the primary visual (Vis) and surrounding areas (Andermann et al., 2011; Marshel et al., 2011; Wang and Burkhalter, 2007), which decreased after whisker training (Figures 2C and S2A; subtraction between Novice and Expert mice images; Wilcoxon rank-sum test, *p*<0.05; for details, see STAR Methods). Stimulation of the C2 whisker evoked two focal responses, in the primary and secondary whisker somatosensory areas (wS1, wS2), both in Novice and Expert mice (Figure 2C). Immediately after, activity transiently decreased in orofacial areas including the primary tongue/jaw sensory and motor areas (tjS1, tjM1), followed by a widespread gradual increase toward the auditory cue initiating in the primary and secondary motor areas for whisker (wM1, wM2) and tongue/jaw (tjM1, ALM), as well as posterior parietal cortex (PPC) and limb/trunk areas. These positive and negative responses during the delay period were selective to Hit trials of Expert mice (Figures 2C, S2B-C, and Videos S3-4). We further quantified response selectivity of different cortical regions for Hit and Correct-rejection trials by comparing their trial-by-trial activity based on ROC analysis (Figure 2D; see STAR Methods). Across all cortical regions tested, selectivity was significantly enhanced in Expert compared to Novice mice during the delay period and response window (*p*<0.05; non-parametric permutation test). Therefore, important learning-induced global changes of information processing emerged during the delay period.

To control for hemodynamic effects of the wide-field fluorescence signal (Makino et al., 2017), we also imaged transgenic mice expressing an activity-independent red fluorescent protein, tdTomato, which has excitation and emission spectra similar to RCaMP (Figure S3; 57 sessions from 7 Expert mice). We imaged RCaMP and tdTomato mice at the same baseline fluorescence intensity (Figure S3E; *p*=0.80, Wilcoxon rank-sum test, n=7 RCaMP mice and n=7 tdTomato mice) by adjusting illumination light power and using identical excitation and emission filters. The tdTomato control mice showed significantly smaller task-related changes in fluorescence than the RCaMP mice (Figure S3A-D; subtraction between RCaMP and tdTomato mice images; Wilcoxon rank-sum test, *p*<0.05). In visual cortex of both RCaMP and tdTomato mice, negative intrinsic signals were evoked around 1 s after the visual stimulus. However, the short whisker stimulation evoked a rapid positive sensory response only in RCaMP mice, and no clear response was evoked in tdTomato mice (Figure S3F). On the other hand, some positive intrinsic optical signals were evoked in midline and frontal regions of tdTomato mice, but the amplitude of these signals was significantly smaller than for RCaMP mice (Wilcoxon rank-sum test, *p*<0.05). These results suggest that the spatiotemporal patterns of fluorescence signals in RCaMP mice largely reflected the calcium activity of the cortex.

### Distinct modification of early and late whisker processing in single neurons

To further investigate learning- and task-related cortical dynamics with higher temporal and spatial resolution, we carried out high-density extracellular recordings (Buzsáki, 2004) from 12 brain regions, with guidance from wide-field calcium imaging (Figures 2 and S2), optical intrinsic imaging and previous literature (Esmaeili and Diamond, 2019; Guo et al., 2014; Harvey et al., 2012; Kyriakatos et al., 2017; Le Merre et al., 2018; Mayrhofer et al., 2019; Sippy et al., 2015; Sreenivasan et al., 2016) including: Vis, wS1, wS2, wM1, wM2, tjM1, ALM, PPC, auditory cortex (Aud), the dorsolateral region of striatum innervated by wS1 (DLS), medial prefrontal cortex (mPFC) and the dorsal part of hippocampal area CA1 (dCA1) (Figures 3A and S4A-C). Two areas were recorded simultaneously during any given session. The precise anatomical location of the recording probes was determined by 3D reconstruction of the probes’ tracks using whole-brain two-photon tomography and registration to the Allen atlas (Figures 3A and S4A-C; for details, see STAR Methods) (Lein et al., 2007; Wang et al., 2020). In total, 4,415 neurons - classified as regular spiking units (RSUs) based on their spike waveform - were recorded in 22 Expert mice, and 1,604 RSUs in 8 Novice mice.

**Figure 3.**
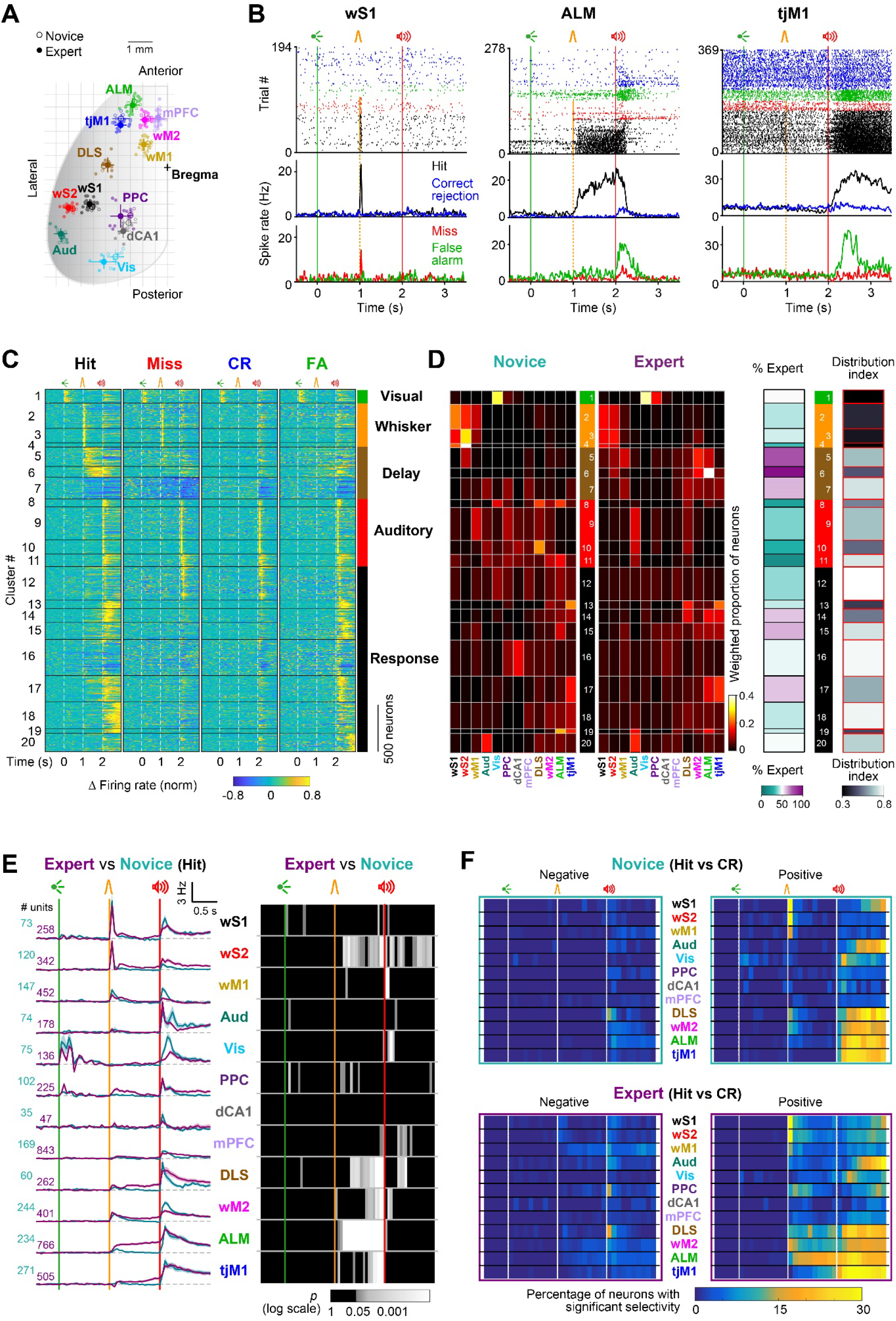
Task epoch-specific processing across single neurons. (A) Reconstructed location of silicon probes registered to the Allen Mouse Brain Atlas in 2D dorsal view in Expert (filled circles) and Novice (open circles). Probes assigned to each anatomical region are shown with different colors and their average coordinates (mean ± SEM) are indicated with larger circles and whiskers. Abbreviations: Medial prefrontal cortex, mPFC; dorsal hippocampal CA1, dCA1; dorsolateral striatum, DLS; other regions as defined in Figure 2B. (B) Example neurons from Expert mice. Raster plots and peristimulus time histograms (PSTHs) for three representative units in wS1, ALM, and tjM1 encoding whisker, delay and licking, respectively. Trials are grouped and colored based on trial outcome. (C) Unsupervised neuronal clustering. Activity maps of all single units from Novice and Expert mice clustered based on their trial-type average normalized firing rate. Black horizontal lines separate different clusters. Labels on the right, indicate the task epoch where the response onset was observed on cluster average response. Only task-modulated clusters (20/24) are shown. (D) Composition of clusters. *Left*, Weighted proportion of neurons within each cluster belonging to different brain regions in Novice and Expert mice. *Right*, Percentage of neurons in each cluster from Novice and Expert mice, and distribution index. To calculate distribution index for each cluster, the probability distribution of the area composition was compared to a uniform distribution and an index between 0 (localized in one area) to 1 (uniformly distributed) was defined. Values are corrected for different sample size in different areas and mouse groups. (E) Population firing rate in Hit trials. *Left*, Baseline-subtracted mean firing rate (mean ± SEM) in each region is superimposed for Expert (purple) and Novice (cyan) mice. *Right, p*-value map of Expert vs Novice mice comparison in 50-ms non-overlapping windows (non-parametric permutation test, FDR-corrected). (F) Proportion of neurons with significant selectivity index in Novice and Expert mice. For individual neurons, selectivity between Hit vs Correct-rejection trials was determined in 100 ms non-overlapping windows based on the area under the ROC curve. Percentage of neurons with significant negative (*left*) or positive (*right*) selectivity in each region is shown across time in Novice (*top*) and Expert (*bottom*) mice. Significance of selectivity was determined using non-parametric permutation tests (*p*<0.05). See also Figures S4-S5 and Videos S5-6.

Single neurons encoded different task aspects such as whisker sensory processing, lick preparation and lick execution (Figure 3B). Assuming that neurons with similar firing dynamics perform similar processing, it is informative to identify those temporal patterns and investigate whether a single pattern is confined or distributed across the brain. We therefore performed unsupervised clustering of neurons according to their temporal firing pattern in different trial types (Hit, Miss, Correct-rejection and False-alarm) by pooling neurons from different brain regions of both Novice and Expert mice (Figures 3C and S5; see STAR Methods). Gaussian mixture model (GMM) clustering (Figure S5A-B) yielded 24 clusters of neurons, among which 20 were modulated in at least one of the task epochs (Hastie et al., 2009). By sorting task-modulated clusters by their onset latency and labeling them based on their task epoch-related response, we analyzed the distribution of clusters across areas along a functional axis (Figure 3C-D). Clusters composed predominantly of neurons from Expert mice were particularly modulated during the Delay period (Clusters 5, 6 and 7) and the Response window (Clusters 14, 15 and 17), and were mainly distributed across different motor-related areas (Figure 3D). Next, we calculated a “distribution index” which quantifies within-area versus between-area composition of clusters (Figures 3D and S5C; for details, see STAR Methods). The distribution index was small for Visual and Whisker clusters, indicating localized distribution of those clusters in specific brain regions. On the other hand, the distribution index was large in the majority of Response clusters, indicating broad distribution of those clusters across brain areas. Across learning, prominent activity patterns remained similar in wS1, wS2 and Vis areas, while it changed in all other regions (Figure S5D).

To reveal spatial changes in neuronal firing following whisker training, we calculated the average time-dependent firing rate for all recording probes (Figure S4D and Videos S5-6) and for the 12 anatomically defined areas (Figure 3E). The visual cue evoked responses localized in Vis and PPC of Novice and Expert mice. Following the auditory cue, excitation rapidly covered all recorded regions in both mice groups. Major changes following whisker training appeared in the delay period between the whisker and auditory stimuli. Similar to deactivation patterns of orofacial cortex revealed by wide-field imaging (Figure 2C), tjM1 showed a transient suppression of firing after whisker stimulation in Expert mice. The whisker stimulus also evoked a widespread excitation across whisker sensorimotor areas (wS1, wS2, wM1, wM2), as well as PPC, DLS and ALM with different latencies. The initial excitation was significantly enhanced in wM2 and ALM (non-parametric permutation test, *p*<0.05). Firing rates of all areas in Novice mice returned to baseline levels shortly after whisker stimulation, whereas in Expert mice wS2, PPC, DLS, wM2, ALM and tjM1 neurons showed increased activity in different parts of the delay. PPC neuronal firing remained elevated only during the first part of the delay period, returning to baseline before the auditory cue, while the activity of wM2, DLS and tjM1 neurons ramped up towards the lick onset. Average neuronal firing in ALM remained elevated throughout the entire delay period. These results suggest that the whisker training enhanced the initial distributed processing of the whisker stimulus, and formed the memory of a licking motor plan among higher-order areas of whisker and tongue/jaw motor cortices, while introducing a transient inhibitory response in tjM1.

We further investigated what was encoded in the acquired neural activity by considering other trial types. First, we found that the pronounced delay period activity during Hit trials was absent in Miss trials, and thus correlated with percept (Figure S4E-F). Second, we quantified the selectivity of single neurons for whisker detection and delayed licking by comparing their trial-by-trial spiking activity in Hit and Correct-rejection trials based on ROC analysis (Figure 3F; see STAR Methods). We found that a significantly larger percentage of neurons became selectively recruited during the delay period in many areas of the Expert mice, suggesting the possible involvement of widespread cortical networks in the acquisition of motor planning for delayed licking (*p*<0.05; non-parametric permutation test).

### Active suppression of orofacial sensorimotor areas

In the delay period, Expert mice showed a transient suppression in broad orofacial sensorimotor cortices selectively in Hit trials (Figures 2, 3, S2 and S4). The suppression of activity in this region coincided with the onset of the whisker-evoked excitation in adjacent secondary motor cortices including ALM (Figure 4A). This inhibition could contribute to suppressing immediate licking in response to the whisker stimulus. To test this hypothesis, we first compared trials in which mice successfully withheld licking until the end of the delay period (Hit), with trials in which mice made premature licking following the whisker stimulus (Early licks). We found that tjM1 activity was significantly suppressed in Hit compared to Early lick trials (Figure 4B), in both calcium imaging signals (tjM1: *p*=0.040; Wilcoxon signed-rank test) and neuronal firing rate (tjM1: *p*=0.017; non-parametric permutation test). Next, to evaluate the causal role of tjM1 in the suppression of premature licking, we optogenetically manipulated tjM1 activity during task execution (Figure 4C). Activation of tjM1 in transgenic mice expressing ChR2 in excitatory neurons (Emx1-ChR2) increased the fraction of Early licks (Figure 4C; n=19 sessions in 6 Expert mice, Light-off versus Light trials, No-Go trials: *p*=4.27×10^−4^, Go trials: *p*=1.94×10^−3^; Wilcoxon signed-rank test). Conversely, inactivation of tjM1 in transgenic mice expressing ChR2 in GABAergic inhibitory neurons (VGAT-ChR2) (Guo et al., 2014) significantly reduced premature licking (Figure 4C; 32 sessions in 9 mice, Light-off versus Light trials, Go trials: Whisker: *p*=0.018, Delay: *p*=2×10^−6^; Wilcoxon signed-rank test). The opposite effect of these optogenetic manipulations indicates that the behavioral changes are not visually-induced by the stimulation light. Altogether, these results suggest that the tjM1 suppression acquired in Expert mice plays an important role in delaying the lick response.

**Figure 4.**
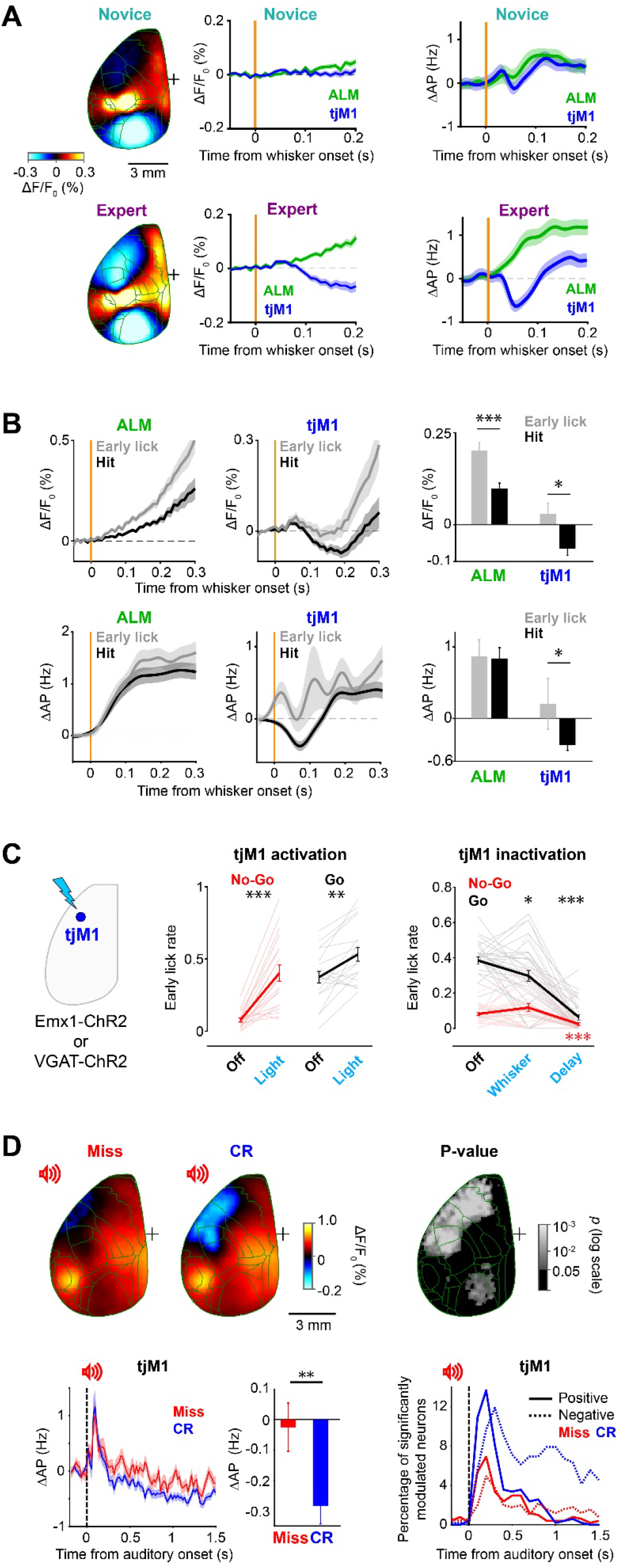
Active suppression of orofacial sensorimotor areas. (A) Transient suppression of tjM1 in Hit trials of Expert mice. Grand average wide-field image (170 ms after whisker onset) of Novice (n=62 sessions, *upper left*) and Expert (n=82 sessions, *lower left*) sessions from 7 mice, calcium traces (*middle*, mean ± SEM) and firing rates (*right*, mean ± SEM) in tjM1 and ALM after whisker stimulus. For the calcium signal, the mean during 50-ms period before whisker stimulation is subtracted, and for spiking data, the mean during 200 ms before whisker onset is subtracted. (B) tjM1 suppression during delayed licking in Expert mice. *Top*, calcium traces averaged (mean ± SEM) across Hit and Early lick trials in ALM (*left*) and tjM1 (*middle*), and comparison of signal amplitude in the suppression window (*right*, 160-210 ms after whisker stimulus; n=82 sessions from 7 mice; ALM: *p*=2.93×10^−4^, tjM1: *p*=0.040; Wilcoxon signed-rank test, FDR-corrected). Mean signal during 50 ms period before whisker onset is subtracted. *Bottom*, average spiking activity (mean ± SEM) in Hit vs Early lick trials in ALM (*left*) and tjM1 (*middle*) and comparison in the suppression window (*right*, 50-100 ms; ALM: n= 766 neurons, *p*=0.466, tjM1: 377 neurons, *p*=0.017, non-parametric permutation test, FDR-corrected). Mean spike rate during 200 ms before whisker stimulus is subtracted. Trials with first lick latency ranging from 300 ms to 1000 ms after whisker stimulus onset were selected for Early lick trials. (C) Causal contribution of tjM1 activity to delayed licking. *Left*, Optogenetic activation and inactivation of tjM1 were performed in Emx1-ChR2 and VGAT-ChR2 transgenic mice, respectively. *Middle*, Fraction of early lick trials in Go and No-Go conditions upon tjM1 activation and no light control trials (n=19 sessions in 6 Expert mice; Light-off versus Light trials, No-Go trials: *p*=4.27×10^−4^, Go trials: *p*=1.94×10^−3^; Wilcoxon signed-rank test, FDR-corrected). *Right*, Fraction of early licks in Go and No-Go trials upon tjM1 inactivation during Whisker or Delay epochs (n=32 sessions in 9 Expert mice; Light-off versus Light trials, No-Go trials: Whisker: *p*=0.239, Delay: *p*=1.2×10^−4^; Go trials: Whisker: *p*=0.018, Delay: *p*=2×10^−6^; Wilcoxon signed-rank test, FDR-corrected). Thick lines show mean ± SEM; lighter lines show individual sessions. For details see Methods. (D) Movement suppression in no-lick trials. *Top*, wide-field images 250 ms after auditory cue in Miss (*left*) and Correct-rejection (*middle*) trials, and *p*-value of comparison (*right;* n=82 Expert sessions from 7 mice; *p*-value of Wilcoxon signed-rank test, FDR-corrected). Mean signal during 50 ms period before auditory onset is subtracted. *Bottom*, baseline-subtracted (200 ms prior to auditory cue) average firing rate (mean ± SEM) of tjM1 neurons in Miss vs Correct-rejection trials (*left*) and the comparison of mean tjM1 spike rate during response window (200-1000 ms window after auditory cue; n=377 neurons; *p*=0.005, non-parametric permutation test); percentage of neurons with positive (solid lines) and negative (dotted lines) modulation in Miss (red) and Correct-rejections (blue) trials during the response period compared to baseline (*right*) (p<0.05; non-parametric permutation test, FDR-corrected).

To further investigate the relationship between reduction of cortical activity and movement suppression, we compared neural activity after the auditory cue between Correct-rejection and Miss trials, as they likely reflect distinct origins of a “no-lick” response (Figure 4D). We found that the calcium signal in orofacial sensorimotor cortices showed significantly stronger suppression in Correct-rejection trials compared to Miss trials (Figure 4D; *p*<0.05; Wilcoxon signed-rank test). Consistently, the spiking activity in tjM1 during the response window revealed a stronger inhibition in Correct-rejection trials (Figure 4D, *p*=0.005; non-parametric permutation test). Moreover, in the same behavioral epoch, a larger proportion of neurons in tjM1 were negatively modulated in Correct-rejection trials (Figure 4D; *p*<0.05; non-parametric permutation test). These results highlight the correlation and causality between the deactivation of orofacial sensorimotor cortex and active suppression of licking.

### Routing of whisker information to frontal cortex

The brief whisker stimulation allowed us to follow the sequence of evoked responses across cortical regions. Frame-by-frame analysis of high-speed calcium imaging data and high-resolution quantification of spiking activity showed that the whisker stimulus evoked the earliest responses in wS1; activity then spread to wS2, wM1, wM2, and finally reached ALM (Figures 5A and S6A-C). This earliest sequence of excitation, as well as the deactivation of tjM1/S1, was significantly enhanced across learning by whisker training (Figure 5A, Wilcoxon rank-sum test, *p*<0.05). This sequential activation and deactivation were diminished when mice failed to lick (Miss trials) (Figure S6D-E; Wilcoxon signed-rank test, *p*<0.05), supporting its involvement in whisker-detection and delayed-licking (see also Figure 4).

**Figure 5.**
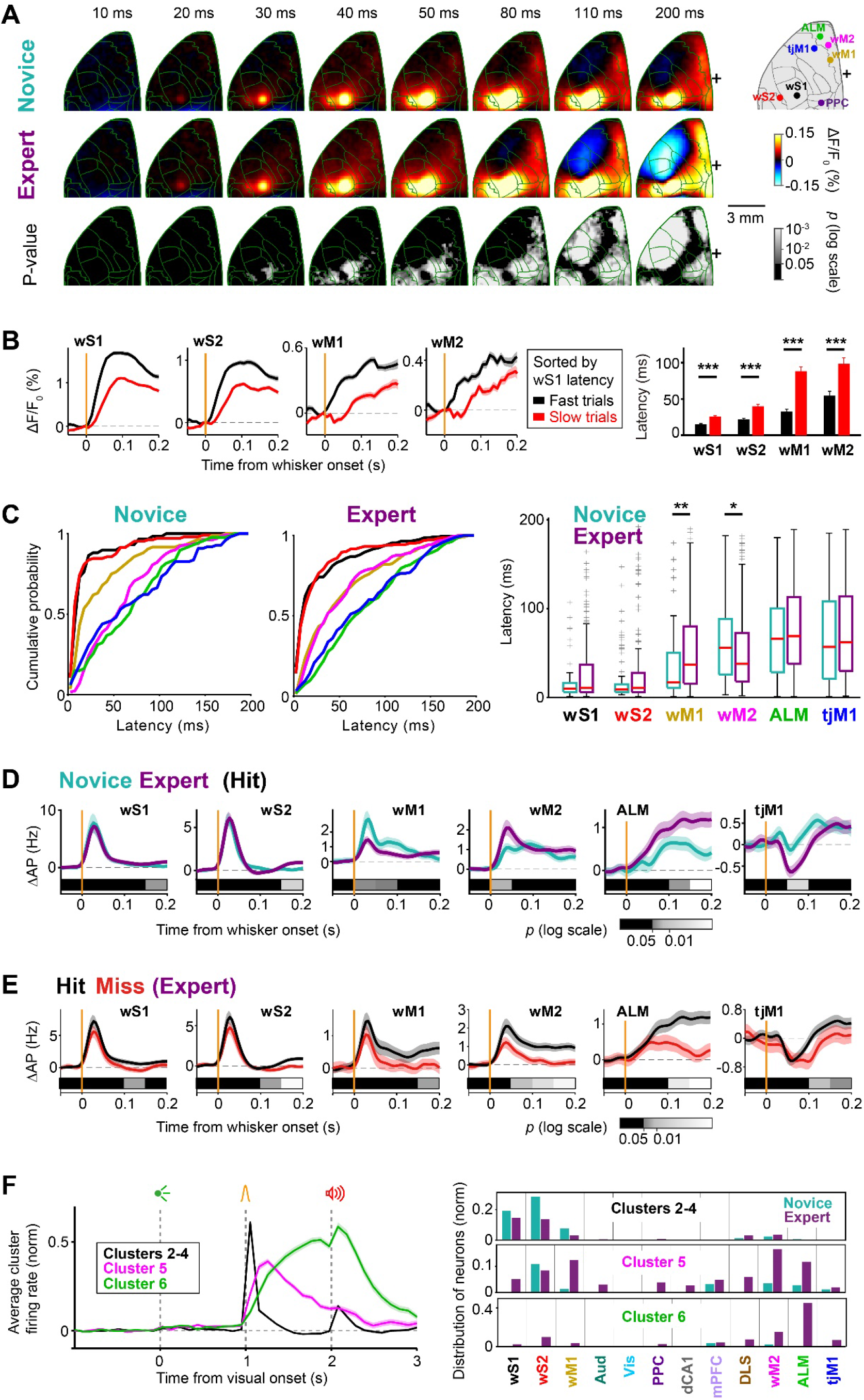
Conversion of a sensory signal into a motor plan. (A) Wide-field signal after whisker stimulus in Novice and Expert mice in Hit trials. Each frame shows the instantaneous calcium activity (10 ms/frame). Mean signal during the 50 ms period before whisker onset is subtracted. From top to bottom, average calcium signal of 62 Novice and 82 Expert sessions from 7 mice, and the statistical significance of the difference (*p*-value of Wilcoxon rank-sum test, FDR-corrected). (B) Propagation of whisker-evoked response latency to downstream regions in Expert mice (82 sessions, 7 mice). *Left*: Calcium traces (mean ± SEM) in different regions were grouped based on single-trial response latencies in wS1. *Right*: Latencies of whisker-evoked calcium response (mean ± SEM) in Fast and Slow trials (wS1: *p*=2.2 ×10^−8^, wS2: *p*=1.1×10^−7^, wM1: *p*=2×10^−9^, wM2: *p*=3.3×10^−4^; Wilcoxon signed-rank test, FDR-corrected). (C) Latency of the whisker-evoked spiking response. Cumulative distribution of single neuron latencies for key cortical areas in Novice (*left*) and Expert (*middle*) mice. Distribution of latencies across different areas and their change across learning (*right*). Boxplots indicate median and interquartile range. Only neurons with significant modulation in the 200 ms window following whisker stimulus compared to a 200 ms window prior to the whisker stimulus are included (*p*<0.05, non-parametric permutation test). Latency was defined at the half maximum (minimum for suppressed neurons) response within the 200 ms window. (D) Early whisker-evoked spiking activity in Hit trials. Baseline-subtracted (200 ms prior to whisker onset) mean ± SEM firing rate across critical cortical areas in Expert and Novice mice are overlaid. Gray horizontal bars represent the *p*-value of Novice/Expert comparison in 50 ms consecutive windows (non-parametric permutation test, FDR-corrected). (E) Spiking activity in Hit vs Miss trials. Baseline-subtracted (200 ms prior to whisker onset) mean ± SEM firing rate across critical cortical areas in Hit and Miss trials of Expert mice are overlaid. Gray horizontal bars represent the *p*-value of Hit/Miss comparison in 50 ms consecutive windows (non-parametric permutation test, FDR-corrected). (F) Whisker and Delay responsive neuronal clusters, related to Figure 3C-D. *Left*, Average normalized firing rate (mean ± SEM) of Whisker (Clusters 2, 3 and 4) and two distinct Delay clusters (Clusters 5 and 6). *Right*, Proportion of neurons within each cluster belonging to different brain regions and groups of mice, related to Figure 3D. See also Figures S5-S6.

To test whether the sequential activation of cortical areas occurs in single trials, we examined whether the variability of the response latency in wS1 propagates to downstream areas in the imaging data. We divided the data into *Slow* and *Fast* trials based on the latency of the whisker-evoked response in wS1 (Figure 5B), and analyzed the latencies in other areas where single-trial analysis of whisker-evoked response latency was feasible (wS2, wM1 and wM2). The latencies of those areas were significantly longer in *Slow* trials (wS1: *p*=2.2×10^−8^, wS2: *p*=1.1×10^−7^, wM1: *p*=2×10^−9^, wM2: *p*=3.3×10^−4^; Wilcoxon signed-rank test), further suggesting a chain of activation from wS1 to the other regions.

We also analyzed the change in response latency in single neuron data between Novice and Expert mice. For neurons with significant firing rate modulation in the 200 ms window following the whisker stimulus compared to the 200 ms before the whisker stimulus (p<0.05, non-parametric permutation test), latency was calculated as the half-maximum (minimum for suppressed neurons) whisker-evoked response (see STAR Methods). The latency of the whisker-evoked response in wM2 was shorter following whisker training, whereas that of wM1 was longer (Figure 5C, wM1: *p*=0.008, wM2: *p*=0.041, Wilcoxon rank-sum test). Moreover, among all areas recorded, wM2 showed the earliest significant increase in firing upon whisker training (Figure 5D, Novice vs Expert: *p*= 0.015, non-parametric permutation test), as well as the earliest significant difference comparing Hit and Miss trials (Figure 5E, Hit vs Miss: *p*=0.010, non-parametric permutation test).

The neuronal clustering revealed 3 main patterns of activity during the delay period (Figures 3C and 5F): i) a fast and transient increase in neuronal activity following the whisker stimulus (Clusters 2-4) that was mostly represented in wS1 and wS2 of both Novice and Expert mice; ii) a slow ramping activity (Cluster 6) that was mostly represented in ALM but only in Expert mice; and iii) the activity of Cluster 5 rose and peaked after Clusters 2-4, but before Cluster 6, and slowly decayed along the delay period, thus bridging the activities of Clusters 2-4 and Cluster 6. Interestingly, Cluster 5 was most prevalent in wM2 of Expert mice, as well as contributing importantly to activity in wS2, wM1 and ALM (Figures 3C-D and 5F).

Altogether (Figure 5C-F), these results highlight the possible role of wM2 as a potential node to bridge sensory processing to motor planning perhaps helping to relay whisker sensory information from wS1/wS2 to ALM.

### Focalized delay period activity in frontal cortex

The most prominent cortical change after whisker training was the emergence of widespread delay period activity (Figures 2 and 3). In the late delay period, Expert mice showed uninstructed, anticipatory movements of whisker, jaw and tongue (Figures 1E-F and S1C-D), which could be broadly correlated with activity across the brain (Musall et al., 2019; Steinmetz et al., 2019). To identify neural activities more directly related to task execution, we leveraged trial-by-trial variability of the neuronal activity and anticipatory movements (Figure 6).

**Figure 6.**
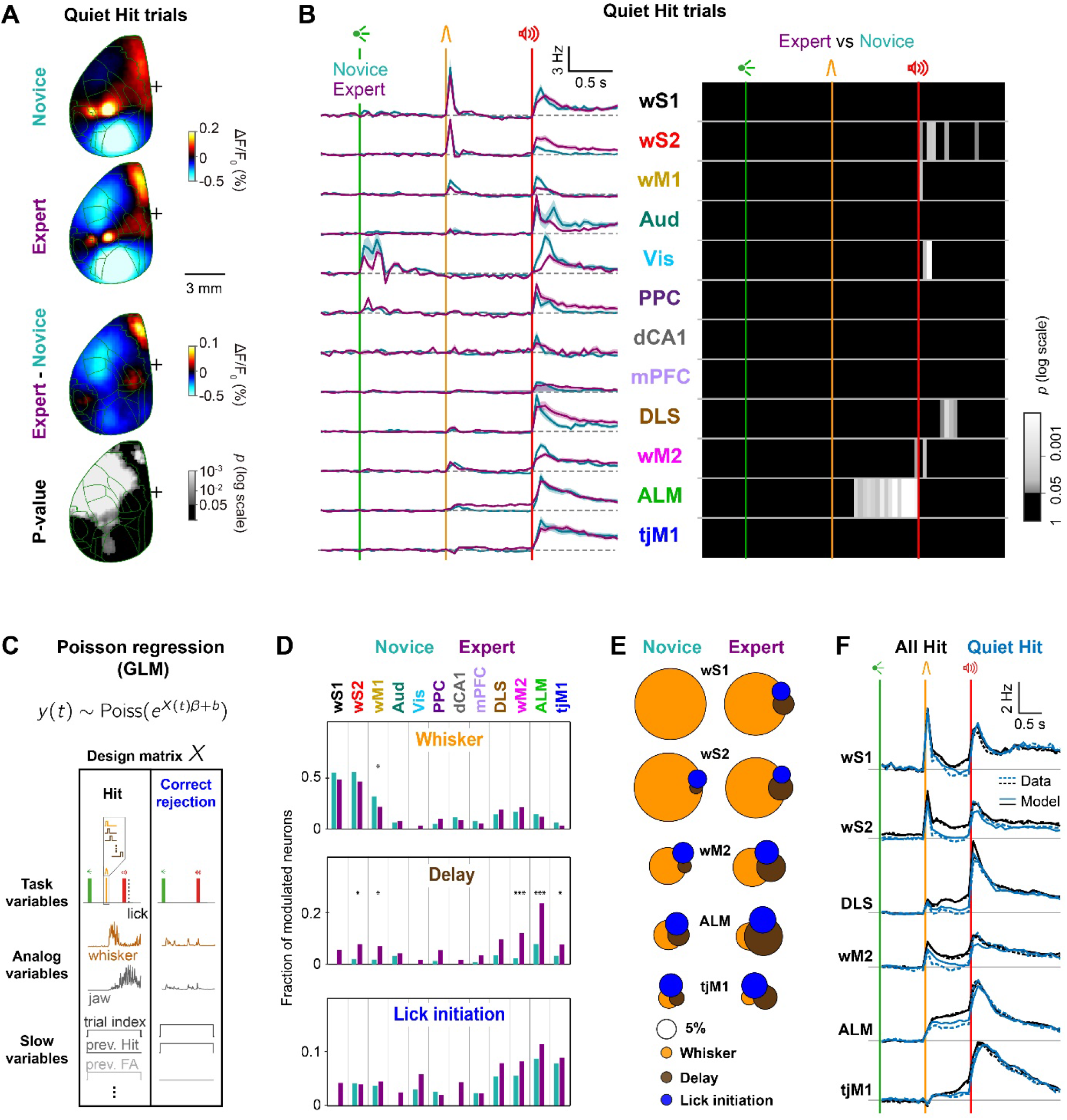
Delay processing beyond preparatory movement. (A and B) Focalized delay activity in Quiet Hit trials. Imaging and neuronal data were averaged across selected Quiet trials with no preparatory jaw movements during the delay period (see STAR Methods). A, Mean wide-field calcium signal in a 50 ms window during the delay period (270 to 320 ms after whisker onset) subtracted by the mean during the 50 ms period before whisker onset. From top to bottom, mean calcium signal of 62 Novice and 82 Expert sessions from 7 mice, their difference, and the statistical significance of the difference (*p*-value of Wilcoxon rank-sum test, FDR-corrected). B, Mean ± SEM firing rate in Expert and Novice mice (*left*) and *p*-value map of Expert/Novice comparison in 50 ms non-overlapping windows (non-parametric permutation test, FDR-corrected) (*right*). (C-F) Poisson encoding model capturing trial-by-trial neuronal variability. C, Schematic of the Poisson encoding model. Concatenated spike trains from Hit and Correct-rejection trials (*y*_*(t)*_) were fitted using a Poisson regression model (GLM). The design matrix (*X*_*(t)*_) included different types of task-related and movement variables (see STAR Methods). D, Fraction of neurons significantly encoding Whisker (*top*), Delay (*middle*) and Lick initiation (*bottom*) (*p*<0.05, likelihood ratio test, See STAR Methods) in different regions. Asterisks represent significant change comparing the fraction of Novice and Expert neurons (proportion test, ***: *p*<0.001, *: *p*<0.05). E, Venn diagrams showing the amount of overlap among neuronal populations in different regions significantly encoding Whisker, Delay and Lick initiation variables. The sizes of the circles are proportional to the fraction of significantly modulated neurons. F, Comparison of empirical (Data, dotted lines) and reconstructed (Model, solid lines) PSTHs for Quiet (blue) and all (black) trials in Expert mice. See also Figure S7 and Table S1.

First, we separated neural activities by selecting trials in which mice did not make jaw movements during the delay period (Quiet trials) (Figures 6A-B, S7A; see STAR Methods). When only Quiet trials were considered, the increased calcium activity during the delay became more localized to ALM (Figure 6A). This focal activation emerged across learning (Wilcoxon rank-sum test, *p<*0.05). Electrophysiology data also demonstrated a consistent localization of the neuronal delay period activity (Figure 6B). In Quiet Hit trials, only ALM population firing remained elevated throughout the delay period and was clearly enhanced by whisker training. In the other regions, the whisker-evoked firing during the delay period returned to baseline, just as in Novice mice. Thus, selecting Quiet trials demonstrated that the essential processing in cortex during the delay period is localized in a focal frontal region that includes ALM.

Assessing the impact of movements considering only Quiet trials highlighted the unique activity pattern of ALM during the delay period. However, Quiet Hits represented a minority of all Hit trials in Expert mice (42 ± 2 %; mean ± SEM). Trials with movements during the delay period may carry richer information about how neuronal activity drives behavior. Therefore, to capture neuronal encoding during single trials, we used a generalized linear model (GLM) (Nelder and Wedderburn, 1972) to fit a Poisson encoding model to spiking data of individual neurons including all correct trials (Park et al., 2014) (Figures 6C-F and S7B-F; see STAR Methods). Three types of model predictors were included (Figure 6C): discrete task events (e.g. sequential boxcars time-locked to sensory stimuli and first lick onset); analog movement signals (whisker, tongue and jaw speed); and slow variables capturing motivational factors (e.g. current trial number) and trial history (e.g. outcome of the previous trial). We assessed fit quality using predictor-spike mutual information and selected only the neurons with a good quality of fit for the rest of analysis (Cover and Thomas, 1991; Gerstner et al., 2014) (Figure S7C; see STAR Methods). The contribution of each model variable to the neuron’s spiking activity was tested by re-fitting the data after excluding the variable of interest (reduced model) and comparing the fit quality to the model including all variables (full model), using a likelihood ratio test (Figures 6D and S7D) (Buse, 1982).

Whisker-related sensorimotor areas (wS1, wS2, wM1 and wM2) had the largest proportion of neurons significantly modulated by whisker stimulus in the first 100 ms, in both Novice and Expert mice (Figures 6D and S7E). The fraction of Whisker encoding neurons decreased across whisker training in wM1 (*p*=0.029). In contrast, Delay encoding neurons - that were significantly modulated between 100 ms and 1 s after the whisker stimulus (Figures 6D and S7E) - were mainly found in ALM, but also in wM2, which was strikingly enhanced by whisker training (*p*=5×10^−5^). Some neurons in wM2, ALM, tjM1 and DLS were found to be significantly modulated during the 200 ms prior to the lick onset, before and after whisker training (Figures 6D and S7E), reflecting the licking initiation signal in these areas beyond those captured by orofacial movements or sound onset predictors in the model.

We next asked to what extent the same neurons encode different task variables. To address this question, we quantified the degree of overlap across populations of Whisker, Delay and Lick initiation encoding neurons in the key areas of interest and visualized it using Venn diagrams (Figure 6E). We found that enhanced Delay and Lick initiation encoding populations were largely non-overlapping. Finally, we asked whether our encoding model, fitted using all trials, can reproduce neuronal activity in Quiet trials (Figures 6F and S7F). Model-reconstructed PSTHs after removing movement-related regressors confirmed that neurons in ALM kept their firing throughout the delay period, while the firing in other areas returned to baseline, in agreement with the empirical data. This result supports the model validity and highlights the prominence of ALM for motor planning.

### Temporally-specific causal contributions of different cortical regions

Imaging and electrophysiology data suggested multiple phases of neural processing for whisker-detection, motor planning and delayed-licking. To examine the causal contribution of cortical regions in each of these phases, we performed spatiotemporally-selective optogenetic inactivation in transgenic mice expressing ChR2 in GABAergic neurons (n=9 VGAT-ChR2 mice). We applied blue light pulses to each brain region through an optical fiber randomly in one-third of the trials, occurring in one of the four temporal windows (Figure 7A): Baseline (from visual cue onset to 100 ms before whisker stimulus onset), Whisker (from 100 ms before to 200 ms after whisker stimulus onset), Delay (from 200 ms to 1000 ms after whisker stimulus onset), or Response (from 0 ms to 1100 ms after auditory cue onset).

**Figure 7.**
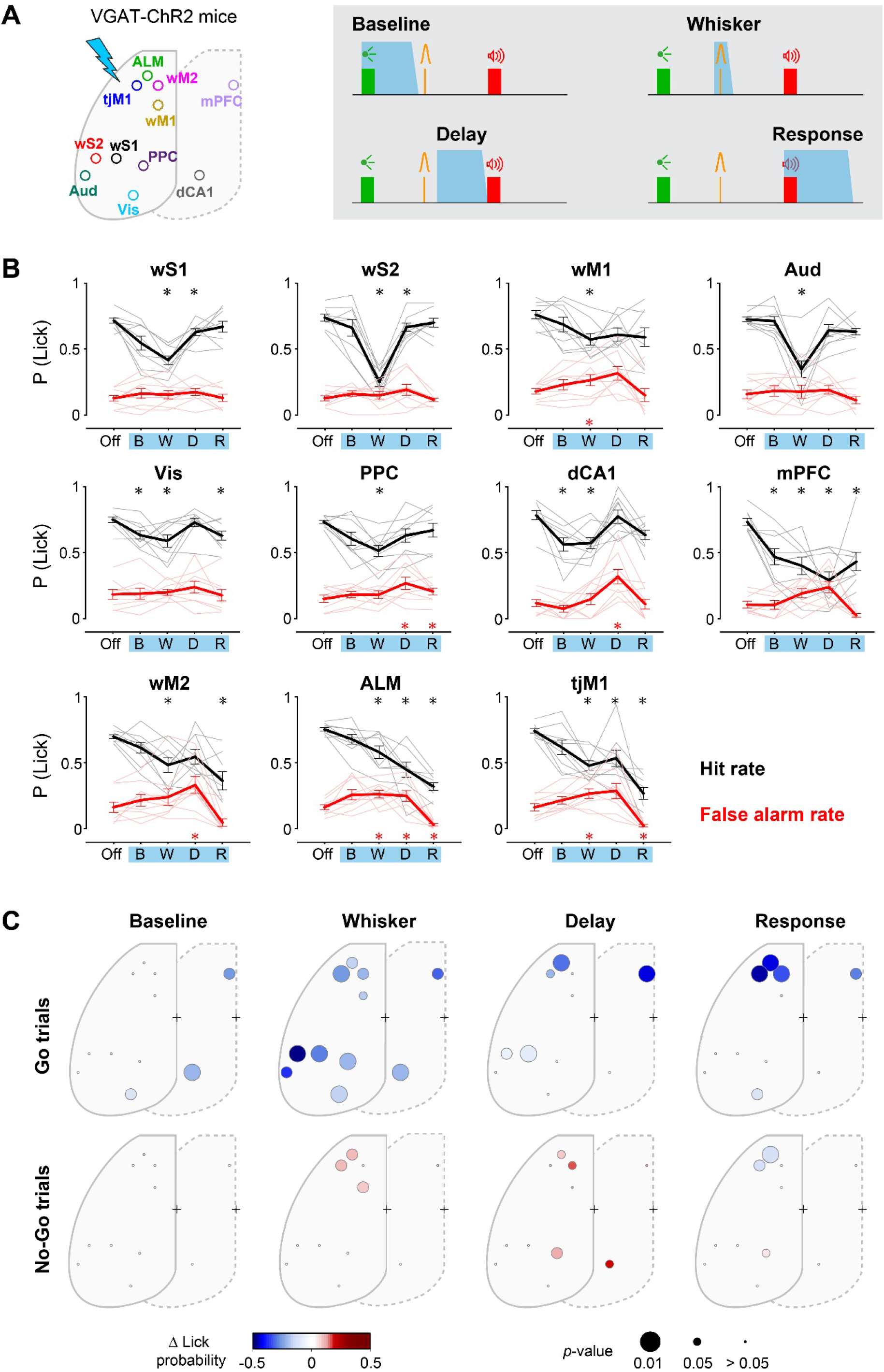
Spatiotemporal causal map of behavioral impact. (A) Spatiotemporally specific optogenetic inactivation in VGAT-ChR2 transgenic mice. Blue shaded areas represent inactivation windows across the trial-timeline. (B) Behavioral impact of optogenetic inactivation across time windows for each brain region (mean ± SEM). For each area, Hit rate (black) and False-alarm rate (red) are plotted for Light-off (Off), Baseline (B), Whisker (W), Delay (D) and Response (R) windows. Asterisks represent significant difference comparing Hit (black) or False-alarm (red) in light trials vs light-off trials (n=9 mice; *, *p*<0.05; Wilcoxon signed-rank test, Bonferroni correction for multiple comparison). (C) Spatiotemporal map of behavioral impact of focal inactivation in Go (*top*) and No-Go trials (*bottom*). Circles represent different cortical regions labeled on the schematic in Panel A; color shows change in Lick probability and circle size shows the *p*-value of the significance test comparing light trials vs light-off trials (n=9 mice, Wilcoxon signed-rank test, Bonferroni correction for multiple comparison). See also Figure S8.

Inactivation in different time windows provided spatiotemporal maps of the behavioral impact (Figures 7B-C and S8). During the Baseline window, a significant decrease in Hit rate occurred after inactivation of Vis, dCA1 and mPFC (Light-off versus Light, Vis: *p*=0.031, dCA1: *p*=0.016, mPFC: *p*=0.031; Wilcoxon signed-rank test). During the Whisker window, a significant decrease in Hit rate occurred in every region tested with the strongest impact in wS2 (Light-off versus Light, *p*=0.016; Wilcoxon signed-rank test). During the Delay period, inactivation of ALM and mPFC produced a strong reduction in Hit rate (Light-off versus Light, ALM: *p*=0.016, mPFC: *p*=0.016; Wilcoxon signed-rank test). Finally, during the Response window, when the licking behavior had to be executed, inactivation of tongue-related tjM1 and ALM, but also whisker-related wM2, impaired behavior by decreasing both Hit and False-alarm rate (Light-off versus Light, tjM1: *p*=0.016, ALM: *p*=0.016, wM2: *p*=0.016; Wilcoxon signed-rank test), supporting the causal involvement of the lick initiation-encoding of wM2 neurons (Figure 6D). The differential impact of inactivating nearby cortical regions is consistent with high spatiotemporal specificity of our optogenetic manipulations. Inactivation during the Whisker and Delay periods also broadly reduced the fraction of premature licking and reduced preparatory movements, with spatiotemporal specificities relatively similar to those observed in Hit rate changes (Figure S8). Thus, spatiotemporal mapping of causal impacts suggests that critical whisker processing is initially distributed across diverse cortical regions, and then converges in frontal regions for planning lick motor output, in agreement with neural activity.

To directly compare the obtained causal maps with observed neural correlations, we quantified the difference in firing rate between Hit versus Correct-rejection and the change in Hit rate upon optogenetic inactivation, for each brain area and time window (Figure 8A). If a brain region is critically involved in task execution, neural activity in that area would code behavioral decision (large Hit-Correct rejection difference), and its inactivation would cause behavioral impairments (strong decrease in Hit rate). This is further quantified by an involvement index as the product of the two terms described above (Figure 8B). The involvement index during the Whisker period was largest in wS2 and wS1 (mean ± SEM, wS2: 0.7 ± 0.11, *p*<0.01, wS1: 0.58 ± 0.11, *p*<0.05; non-parametric permutation test versus other areas) highlighting these areas as the main nodes of whisker sensory processing. During the Delay period, ALM had the largest involvement index (mean ± SEM, ALM: 0.48 ± 0.09, p<0.001; non-parametric permutation test versus other areas). Although, mPFC inactivation during the delay provoked the largest reductions in hit rate, there was little change in neuronal activity in this area, resulting in small involvement values. The most critical areas in the Response window were tjM1 and ALM (mean ± SEM, tjM1: 1.16 ± 0.15, *p*<3×10^−5^, ALM: 0.76 ± 0.09, *p*<0.05; non-parametric permutation test versus other areas). This reflects the prominent role of tjM1 in licking execution. Interestingly, wM2 had a moderate but significant involvement index in all three time-windows, supporting its possible role in bridging sensory processing and motor execution.

**Figure 8.**
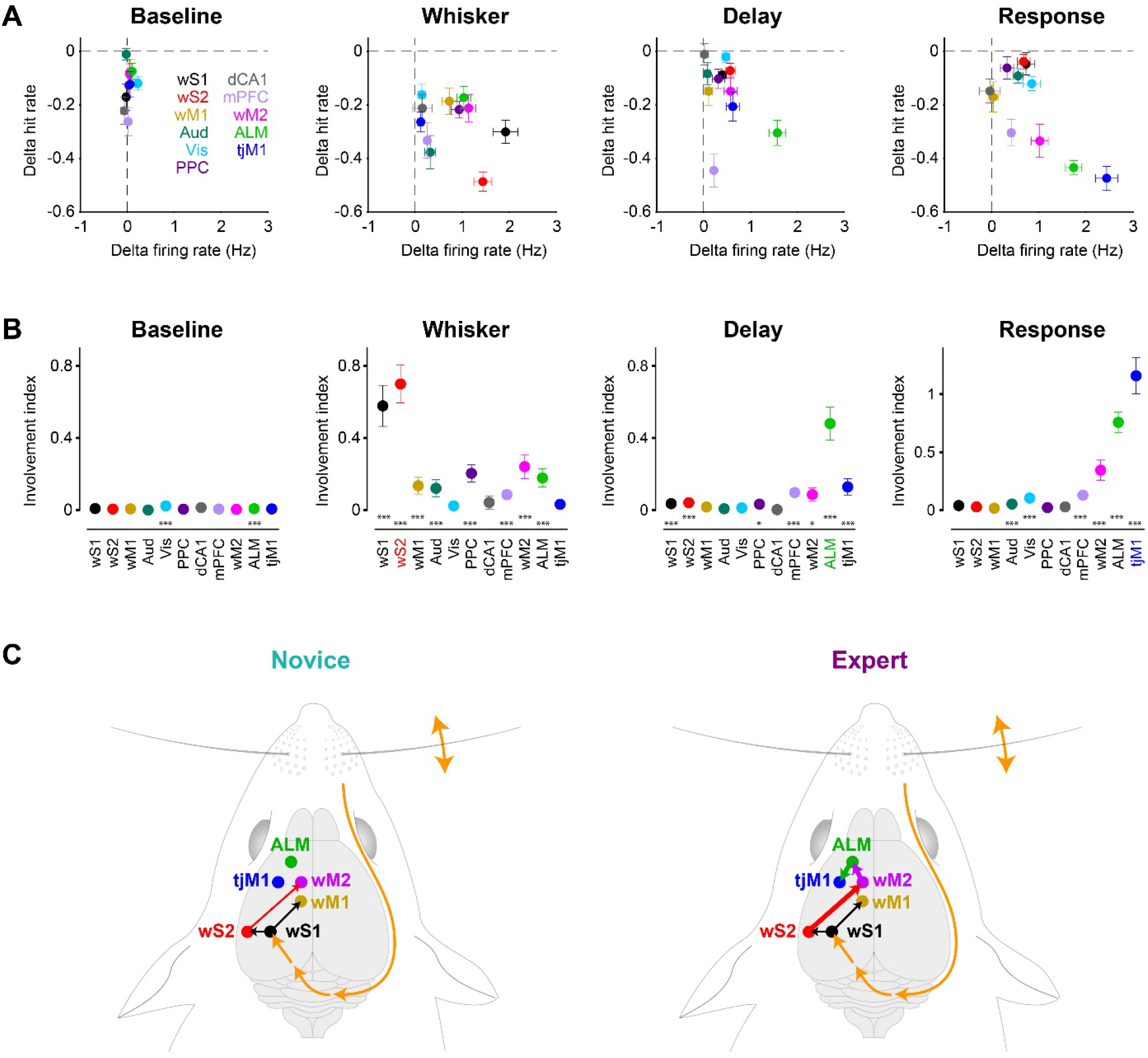
Task-epoch specific involvement of cortical regions. (A) Trial specific neural firing versus behavioral impact of inactivation. Firing rate difference in Hit vs Correct-rejection trials (n=22, Expert mice) in each time window is plotted against the change in hit rate upon optogenetic inactivation (n=9, VGAT-ChR2 mice). Temporal windows are defined similarly to Figure 7. Circles represent mean values in different areas; horizontal and vertical error bars show SEM. (B) A causal involvement index was defined as the region- and epoch-specific absolute value of the difference in firing rate comparing Hit and Correct-rejection trials (n=22, Expert mice) multiplied by the change in hit rate induced by optogenetic inactivation (n=9, VGAT-ChR2 mice). Error bars are obtained from bootstrap (see STAR Methods) and represent standard deviation (bootstrap standard error). Asterisks represent significance level (*, *p*<0.05; ***, *p*<0.001; non-parametric permutation test, Bonferroni correction for multiple comparison). (C) Proposed cortical circuits connecting whisker somatosensory cortex to tongue/jaw motor cortex upon task learning.

## DISCUSSION

We found converging evidence for the temporally distinct involvement of diverse cortical regions in delayed sensorimotor transformation using an array of complementary technical approaches. Our analyses of the learning-induced changes in causal neural activity revealed three key findings further discussed below: i) widespread neuronal delay-period activity was dominated by preparatory movements, but essential causal neuronal delay-period activity was predominantly localized to ALM; ii) sequential activation of cortical regions wS1, wS2, wM2 and ALM suggests the possible contribution of a corticocortical pathway for whisker sensory information to reach ALM, with wM2 showing the earliest increase in sensory-evoked response across learning; and iii) suppression of orofacial sensorimotor cortex in the early delay period, likely contributing to inhibition of premature licking.

### Essential cortical delay period activity in ALM

Broad regions of cortex showed elevated activity in Expert mice during the delay period in Hit trials (Figures 2 and 3), correlating with preparatory movements (Figures 1 and 6). These results are thus in good agreement with widespread motor-related cortical activity (Musall et al., 2019; Steinmetz et al., 2019; Stringer et al., 2019). When we analyzed only trials free from the delay period preparatory movements, wide-field imaging and electrophysiology demonstrated a localized excitatory activity in a small region of secondary motor cortex including ALM (Figure 6A-B). Inactivation of ALM during the delay period was highly effective in reducing hit rates in the subsequent Response period (Figure 7). Essential causal neuronal delay period activity therefore appears to be predominantly localized to ALM (Figure 8A-B), in good agreement with previous closely-related tasks (Guo et al., 2014; Li et al., 2015).

By accounting for movement contributions using linear regression analysis of trial-by-trial variability, we found that most delay period-responsive neurons were indeed localized in ALM, but that the fraction of delay encoding neurons was also significantly enhanced by learning in wS2, wM1, wM2 and tjM1 (Figure 6C-E). Furthermore, during the delay period, inactivation of several cortical areas including not only ALM but also wS1, wS2, mPFC and tjM1 significantly reduced hit rates (Figure 7). Indeed, causal contributions to the Delay period measured by the Involvement Index were also significant in wS1, wS2, PPC, mPFC, wM2 and tjM1, as well as ALM. In addition to the strongest causal involvement found for ALM, these causal impacts observed in broader cortical areas during delay period might in part result from reduced preparatory movements induced by inactivation (Figure S8B-C). The preparatory movements, which were most prominent in Hit trials of Expert mice, may thus contribute a form of embodied sensorimotor memory in which ongoing movements might help maintain a plan for delayed licking (Mayrhofer et al., 2019).

During the Delay period, mPFC inactivation had the largest impact on hit rate across the tested areas (Figure 7). However, we did not find robust sustained activity in mPFC during this window for maintenance of the motor plan. Interestingly, mPFC inactivation during all task epochs (including baseline) impaired behavior. One possibility is that the observed behavioral effect relates to the representation of task rules (Durstewitz et al., 2010), behavioral strategy (Powell and Redish, 2016) or motivation (Popescu et al., 2016).

### A putative corticocortical signaling pathway linking sensory to motor cortex through learning

Our measurements at high spatiotemporal resolution revealed a rapid sequential activation of cortical areas evoked by whisker-deflection, ultimately reaching ALM in Hit trials of Expert mice. The earliest cortical response to whisker stimulus occurred in wS1 and wS2, which changed relatively little after whisker training (Figures 2, 3 and 5). This initial processing was essential as shown by optogenetic inactivation (Figure 7), and therefore wS1 and wS2 appear to form the cortical starting points for task execution, in agreement with previous studies of whisker detection tasks without a delay period (Kwon et al., 2016; Kyriakatos et al., 2017; Le Merre et al., 2018; Mayrhofer et al., 2019; Miyashita and Feldman, 2013; Sachidhanandam et al., 2013; Yang et al., 2016).

Sensory cortical areas project directly and strongly to frontal cortex through parallel pathways, with wS1 innervating wM1, and wS2 innervating wM2 (Ferezou et al., 2007; Mao et al., 2011; Oh et al., 2014; Sreenivasan et al., 2017). Whisker-deflection evoked rapid sensory responses in these downstream motor regions. Interestingly, the sensory response in wM2 showed the earliest significant increase in whisker-evoked firing and a decrease in response latency across learning (Figure 5C-D), whereas a decrease in amplitude and increase in latency was found in wM1. Neuronal activity in wM2 also showed the earliest choice-related activity when comparing Hit and Miss trials (Figure 5E). Thus, wM2 might serve as a key node in the corticocortical network to begin the process of converting a whisker sensory stimulus into longer-lasting preparatory neuronal activity. Shortly after wM2 activation, ALM, an important premotor area for control of licking (Guo et al., 2014; Li et al., 2015; Mayrhofer et al., 2019), started to increase firing (Figure 5). Through cortico-cortical connectivity (Luo et al., 2019), activity in wM2 could contribute directly to exciting its neighbor region ALM, which manifested the most prominent delay period activity through whisker training (Figures 3 and 6), consistent with previous studies (Chen et al., 2017; Li et al., 2015).

Our results suggest a hypothesis for a minimal cortical network connecting whisker sensory coding to preparatory neuronal activity for motor planning: a pathway wS1 → wS2 → wM2 → ALM could be the main stream of signal processing (Figure 8C). Some of the most prominent whisker-related changes through whisker training occurred in wM2 and ALM, and it is possible that reward-related potentiation of synaptic transmission between wS2 → wM2 and wM2 → ALM could underlie important aspects of the present learning paradigm. All of these cortical areas are likely to be connected through reciprocal excitatory long-range axonal projections, which could give rise to recurrent excitation helping to prolong firing rates of neurons in relevant brain regions during the delay period of Hit trials. Interestingly, in a related whisker detection task without a delay period, enhanced reciprocal signaling between wS1 and wS2 has already been proposed to play an important role (Kwon et al., 2016; Yamashita and Petersen, 2016). It is also important to note that a large number of subcortical structures are also likely to be involved in task learning and performance including thalamus (El-Boustani et al., 2020; Guo et al., 2017), basal ganglia (Sippy et al., 2015) and cerebellum (Chabrol et al., 2019; Gao et al., 2018).

### Lick and No-Lick signals in tjM1

In Expert mice, we found that the whisker stimulus evoked a sharp deactivation broadly across orofacial sensorimotor cortex, including tjM1, an area thought to be involved in the initiation and control of licking (Mayrhofer et al., 2019). In contrast, tjM1 neurons were activated soon after whisker deflection in a previous study of a detection task without a delay period before licking (Mayrhofer et al., 2019). One interesting possibility is thus that the deactivation in tjM1 develops through learning of a task where suppression of immediate licking is demanded. In support of this hypothesis, here, we found that premature early licking during the delay period was accompanied by reduced suppression of tjM1 (Figure 4B), and that activation of tjM1 increased early licks whereas inactivation of tjM1 reduced early licks (Figure 4C). We furthermore found that tjM1 activity was suppressed after the auditory cue in Correct-rejection trials where mice are supposed to suppress licking, compared to Miss trials where mice failed to lick, suggesting that the reduction of activity in orofacial cortex reflects active response inhibition (Figure 4D). Finally, inactivation of tjM1 in the Response window evoked the strongest decrease in hit rates further supporting the causal involvement of this area in the control of licking (Figure 7).

Previous studies in human subjects have suggested the importance of inhibitory mechanisms for preventing actions from being emitted inappropriately (Chikazoe et al., 2009; Duque et al., 2017). Parallel suppression and activation during a delay period might be a common principle of response preparation preserved across species (Cohen et al., 2010). Here, we reveal causal contributions of inhibitory and excitatory cortical delay period activity in a precisely-defined task, and, as a hypothesis, we put forward a specific corticocortical circuit which could contribute to task learning and execution, requiring future further experimental testing.

## Supporting information

Video S1 WF Novice Hit

Video S2 WF Expert Hit

Video S3 WF Novice Miss

Video S4 WF Expert Miss

Video S5 AP Novice Hit

Video S6 AP Expert Hit

## Acknowledgments

We thank Fritjof Helmchen for TIGRE1.0-RCaMP mice, Eloise Charrière and Candice Stoudmann for management of mouse colonies and help with behavioral training, Sai Krishna Teja Sadhu for help with image analysis, and Flavio Martinelli for help with video-tracking. This work was supported by the Swiss National Science Foundation (310030B_166595, 31003A_182010 and CRSII5_177237 to CCHP, 200020_165538 to SPM) and the European Research Council (ERC-2011-ADG 293660) to CCHP, European Union’s Marie Skłodowska-Curie Actions (665667, 798617), the Research Foundation for Opto-science and Technology, the Brain Science Foundation, the Japan Society for the Promotion of Sciences, and the Ichiro Kanehara Foundation to KT.

## Author Contributions

VE, KT, SC and CCHP conceptualized the study; VE and KT developed neural and behavioral experiment setups; VE, KT, and MB obtained neural and behavioral data; VE, KT, ABL and AO obtained histological data; GF and YL built the two-photon tomography system; VE, KT, SPM and AM analyzed the data; WG advised clustering and fitting of neuronal data; VE, KT, SC and CCHP wrote the manuscript; all authors discussed and edited the manuscript; and CCHP provided overall supervision.

## Declaration of Interests

C.C.H.P. is a member of the advisory board of *Neuron*.

## STAR METHODS

## RESOURCE AVAILABILITY

### Lead contact

Further information and requests for resources and reagents should be directed to and will be fulfilled by the Lead Contact, Carl Petersen.

### Materials availability

This study did not generate new unique reagents.

### Data and code availability

The complete data set and Matlab analysis code are freely available at the open access CERN Zenodo database https://doi.org/10.5281/zenodo.4720013.

## EXPERIMENTAL MODEL AND SUBJECT DETAILS

All procedures were approved by Swiss Federal Veterinary Office (License number VD-1628) and were conducted in accordance with the Swiss guidelines for the use of research animals. For calcium imaging, we produced RCaMP mice by crossing Emx1-IRES-Cre mice [B6.129S2-Emx1<tm1(cre)Krj>/J, JAX: 005628] (Gorski et al., 2002), CaMK2-tTA mice [B6.Cg-Tg(Camk2a-tTA)1Mmay/DboJ, JAX: 007004] (Mayford et al., 1996), and TITL-R-CaMP mice [TIGRE1.0-RCaMP, B6.Cg-Igs7<tm143.1(tetO-RCaMP1.07)Hze>/J, JAX: 030217, kind gift from Fritjof Helmchen (University of Zurich)] (Bethge et al., 2017). For control imaging, we produced tdTomato mice by crossing VIP-IRES-Cre mice [STOCK Vip<tm1(cre)Zjh>/J, JAX: 010908] (Taniguchi et al., 2011) and LSL-tdTomato mice [B6.Cg-Gt(ROSA)26Sor<tm9(CAG-tdTomato)Hze>/J, JAX: 007909] (Madisen et al., 2010). For optogenetic activation, we produced Emx1-ChR2 mice by crossing Emx1-IRES-Cre mice, LSL-ChR2(H134R)-EYFP mice [B6;129S-Gt(ROSA)26Sor<tm32(CAG-COP4*H134R/EYFP)Hze>/J] (Madisen et al., 2012) and RCaMP mice. For optogenetic inactivation, we used VGAT-ChR2 mice [B6.Cg-Tg(Slc32a1-COP4*H134R/EYFP)8Gfng/J, JAX: 014548] (Zhao et al., 2011). For electrophysiological recording, we used C57BL/6 wild type mice, and VGAT-ChR2 mice, as well as A2A-Cre mice [B6.FVB(Cg)-Tg(Adora2a-cre)KG139Gsat/Mmucd, MMRRC: 036158] (Gong et al., 2007) crossed with LSL-tdTomato mice. Adult male and female mice were at least 6 weeks old at the time of head-post implantation (see below). Mice were kept in a reverse light/dark cycle (light 7 p.m. to 7 a.m.), in ventilated cages at a temperature of 22 ± 2°C with food available ad libitum. Water was restricted to 1 ml a day during behavioral training with at least 2 days of free-access to water in the cage every 2 weeks. All mice were weighed and inspected daily during behavioral training.

## METHOD DETAILS

### Experimental design

This study did not involve randomization or blinding. We did not estimate sample-size before carrying out the study. However, the sample-size in this study is comparable with those used in related studies (Allen et al., 2017; Guo et al., 2014; Harvey et al., 2012; Hattori et al., 2019; MacDowell and Buschman, 2020; Pinto et al., 2019).

### Implantation of metal headpost

Mice were deeply anesthetized with isoflurane (3% with O_2_) and then were maintained under anesthesia using a mixture of ketamine and xylazine injected intraperitoneally (ketamine: 125 mg/kg, xylazine: 10 mg/kg). Carprofen was injected intraperitoneally (100 µl at 0.5 mg/ml) for analgesia before the start of surgery. Body temperature was kept at 37°C throughout the surgery with a heating pad. An ocular ointment (VITA-POS, Pharma Medica AG, Switzerland) was applied over the eyes to prevent them from drying. As local analgesic, a mix of lidocaine and bupivacaine was injected below the scalp before any surgical intervention. A povidone-iodine solution (Betadine, Mundipharma Medical Company, Bermuda) was used for skin disinfection. To expose the skull, a part of the scalp was removed with surgical scissors. The periosteal tissue was removed with cotton buds and a scalpel blade. After disinfection with Betadine and rinsing with Ringer solution, the skull was dried well with cotton buds. A thin layer of super glue (Loctite super glue 401, Henkel, Germany) was then applied across the dorsal part of the skull and a custom-made head fixation implant was glued to the right hemisphere without a tilt and parallel to the midline. A second thin layer of the glue was applied homogeneously on the left hemisphere. After the glue had dried, the head implant was further secured with self-curing denture acrylic (Paladur, Kulzer, Germany; Ortho-Jet, LANG, USA). For electrophysiological recordings a chamber was made by building a wall with denture acrylic along the edge of the bone covering the left hemisphere. Particular care was taken to ensure that the left hemisphere of the dorsal cortex was free of denture acrylic and only covered by super glue for optical access. This intact, transparent skull preparation was used to perform wide-field calcium imaging as well as intrinsic optical signal imaging (IOS) experiments. Mice were returned to their home cages and ibuprofen (Algifor Dolo Junior, VERFORA SA, Switzerland) was added to the drinking water for three days after surgery.

### Skull preparation and craniotomies

For wide-field calcium imaging and optogenetic activation, an intact transparent skull was used as described above. For electrophysiological recordings, up to 10 small craniotomies were made over the regions of interest using a dental drill under isoflurane anesthesia (2-3% in O_2_). The craniotomies were protected using a silicon elastomer (Kwik-Cast, World Precision Instruments, Sarasota, FL, USA). Regions of interest were selected based on the hotspots of activity from wide-field calcium imaging experiments, functionally relevant areas based on previous studies (Esmaeili and Diamond, 2019; Guo et al., 2014; Harvey et al., 2012; Le Merre et al., 2018; Mayrhofer et al., 2019; Sachidhanandam et al., 2013; Sippy et al., 2015; Sreenivasan et al., 2016) and IOS imaging (Lefort et al., 2009). IOS was performed under isoflurane anesthesia (1-1.5% with O_2_) to map the C2-whisker representation in primary and secondary whisker somatosensory cortex (wS1 and wS2), as well as the auditory area (Aud). A piezoelectric actuator was used to vibrate the right C2 whisker, or to generate rattle sounds. Increase in absorption of red light (625 nm) upon sensory stimulation indicated the functional location of the corresponding sensory cortex. For the other regions stereotaxic coordinates relative to bregma were used: primary and secondary whisker motor cortices (wM1: AP 1.0 mm; Lat 1.0 mm and wM2: AP 2.0 mm; Lat 1.0 mm), primary and secondary tongue/jaw motor cortices (tjM1: AP 2.0 mm; Lat 2.0 mm and ALM: AP 2.5 mm; Lat 1.5 mm), visual cortex (Vis: AP -3.8 mm; Lat 2.5 mm), posterior parietal cortex (PPC: AP -2 mm; Lat 1.75 mm), medial prefrontal cortex (mPFC: AP 2 mm; Lat 0.5 mm), dorsal part of the CA1 region of the hippocampus (dCA1: AP -2.7 mm; Lat 2.0 mm) and dorsolateral striatum (DLS: AP 0.0 mm; Lat 3.5 mm). For optogenetic inactivation experiments the bone over the regions of interest was thinned and a thin layer of superglue was applied to protect the skull for stable optical access over days. For the inactivation of mPFC and dCA1 a small craniotomy was made for the insertion of an optical fiber or an optrode.

### Behavioral paradigm

A total of 55 mice were examined in the delayed whisker detection task including 9 RCaMP, 24 wild-type or negative, 6 Emx1-ChR2, 9 VGAT-ChR2 and 7 tdTomato mice. During the behavioral experiments, all whiskers were trimmed except for the C2 whiskers on both sides, and the mice were water restricted to 1 ml of water/day. Mice were trained daily with one session/day and their weight and general health status were carefully monitored using a score sheet. Both groups of mice (Expert and Novice) went through a Pretraining phase which consisted of trials with visual and auditory cues (without any whisker stimulus) (Figure 1C). Mice were rewarded by licking a spout, placed on their right side, in a 1-second response window after the auditory cue onset. Trials were separated 6-8 seconds and started after a quiet period of 2-3 seconds in which mice did not lick the spout. Each trial consisted of a visual cue (200 ms, green LED) and an auditory cue (200 ms, 10 kHz tone of 9 dB added on top of the continuous background white noise of 80 dB). The stimuli were separated with a delay period which gradually was increased to 2 seconds over Pretraining days. Licking before the response period (Early lick) aborted the trial and introduced a 3-5 second timeout. After 3-6 days of Pretraining, mice learned to lick the spout by detecting the auditory cue and to suppress early licking.

The wide-field imaging and electrophysiological recordings from the Novice group of mice was performed when mice finished the Pretraining phase and were introduced to the whisker delay task (Figure 1C). In this phase a whisker stimulus (10 ms cosine 100 Hz pulse through a glass tube attached to a piezoelectric driver) was delivered to the right C2 whisker 1 second after the visual cue onset in half of the trials. Importantly, the reward was available only in trials with the whisker stimulus (Go trials), and time-out punishment (together with an auditory buzz tone) was given when mice licked in trials without the whisker stimulus (No-Go trials) (Figure 1B). Thus, mice were requested to use the whisker stimulus to change their lick/no-lick behavior. Since the whisker stimulus was weak, Novice mice continued licking in most of Go and No-Go trials irrespective of the whisker stimulus and did not show any sign of whisker learning (Figure 1D and S1B).

The Expert mice entered a Whisker-training phase of 2-29 days during which a stronger whisker stimulus (larger amplitude and/or train of pulses) and shorter delays (for some mice) was introduced (Figure S1A). As the mice learned to lick correctly, the whisker stimulus amplitude was gradually returned to a smaller amplitude and delay was extended to 1 second, eventually matching the conditions in Novice mice. Expert mice decreased licking in No-Go trials but increased their premature early licks after the whisker stimulus, as monitored by the piezoelectric lick sensor (Figure 1D, see below). Behavioral hardware control and data collection were carried out using data acquisition boards (National Instruments, USA) and custom-written Matlab codes (MathWorks).

### Quantification of orofacial movements

Contacts of the tongue with the reward spout were detected by a piezo-electric sensor. Continuous movements of the left C2 whisker, tongue and jaw were filmed by a high-speed camera (CL 600 × 2/M, Optronis, Germany; 200 or 500 Hz frame rate, 0.5-or 1 ms exposure, and 512×512-pixel resolution) under blue light or infrared illumination. Movements of each body part were tracked using custom-written Matlab codes. For the imaging sessions, arc regions-of-interest were defined around the basal points for both the whisker and jaw (Mayrhofer et al., 2019). Crossing points on these arcs were detected for the whisker (the pixels with the minimum intensity) and the jaw (pixels with the maximum slope of intensity). A vector was then defined for each pair of basal point and the cross point, and the absolute angle was calculated for each vector with respect to midline. For the electrophysiology sessions, whisker angular position was quantified in a similar manner while movements of tongue and jaw were quantified as the changes in mean image intensity within a rectangular regions-of-interest (ROI) defined separately on the tracks of tongue and jaw. These signals were then normalized to the area covered by tongue and jaw ROIs. Absolute derivatives of orofacial time series (whisker/jaw/tongue speed) were calculated to derive angular whisker speed and normalized tongue/jaw speed.

### Wide-field calcium imaging

Mice were mounted with a 24-degree tilt along the rostro-caudal axis. The red fluorescent calcium indicator R-CaMP1.07 or the red fluorescent protein tdTomato were excited with 563-nm light (567-nm LED, SP-01-L1, Luxeon, Canada; 563/9-nm band pass filter, 563/9 BrightLine HC, Semrock, USA) and red emission light was detected through a band pass filter (645/110 ET Bandpass, Semrock). A dichroic mirror (Beamsplitter T 588 LPXR, Chroma, USA) was used to separate excitation and emission light. Through a face-to-face tandem objective (Nikkor 50 mm f/1.2, Nikon, Japan; 50 mm video lens, Navitar, USA) connected to a 16-bit monochromatic sCMOS camera (ORCA FLASH4.0v3, Hamamatsu Photonics, Japan), images of the left dorsal hemisphere were acquired with a resolution of 256×320-pixels (4×4 binning) aligned in rostro-caudal axis at a frame rate of 100 Hz (10 ms exposure). Behavioral task and imaging were synchronized by triggering acquisition of each image frame by digital pulses sent by the computer for behavioral task control. For each trial, 600 frames (6 seconds) of images were acquired from 1 second before the visual cue onset to 3 seconds after the auditory cue onset. To control for calcium-independent changes in cortical fluorescence (Makino et al., 2017), we imaged transgenic mice expressing tdTomato in vasoactive intestinal peptide-expressing neurons (tdTomato mice) by using the same optical filters as the imaging of RCaMP. tdTomato had excitation and emission spectra similar to RCaMP, and the illumination condition was adjusted so that tdTomato mice and RCaMP mice had comparable fluorescence intensity.

### Electrophysiological recording

Extracellular spikes were recorded using single-shank silicon probes (A1×32-Poly2-10mm-50 s-177, NeuroNexus, MI, USA) with 32 recording sites covering 775 µm of the cortical depth. In each session two probes were inserted in two different brain targets acutely. Probes were coated with DiI (1,1’-Dioctadecyl-3,3,3’,3’-Tetramethylindocarbocyanine Perchlorate, Invitrogen, USA) for post-hoc recovery of the recording location (see below). The neural data were filtered between 0.3 Hz and 7.5 kHz and amplified using a digital headstage (CerePlex™ M32, Blackrock Microsystems, UT, USA). The headstage digitized the data with a sampling frequency of 30 kHz. The digitized signal was transferred to our data acquisition system (CerePlex™ Direct, Blackrock Microsystems, UT, USA) and stored on an internal HDD of the host PC for offline analysis.

### Optogenetic manipulations

Optogenetic activation of tjM1 was performed in 6 Expert Emx1-ChR2 mice with the same transparent skull preparation and 24-deg tilt as the wide-field imaging. 473-nm laser beam (S1FC473MM, Thorlabs) was steered on the cortex by a pair of Galvo mirrors (GVS202, Thorlabs) (Mayrhofer et al., 2019) connected to the wide-field imaging system via a short-pass beam splitter (F38-496SG, Semrock). In a random half of Go and No-Go trials, a single brief laser pulse (duration, 2 ms; diameter, ∼ 400 μm; power, 1.5, 3, 4.5, 6, or 9 mW, randomly selected) was delivered to the tjM1 (2.6 mm lateral and 1.8 mm anterior to the bregma in 24-deg tilt) (Mayrhofer et al., 2019) at 1050 ms after the visual stimulus onset, at which time the neuronal firing in tjM1 showed the maximal suppression in Hit trials. In the other half of trials, the laser pulse with the same parameters was delivered to the edge of the implant as a control stimulation so that mice could not discriminate tjM1-stimulated and non-stimulated trials by visual cues. Both spontaneous and optogenetically evoked Early licks led to trial abortion with time-out, thus preventing any reinforcement of Early licks.

Optogenetic inactivations were performed in 9 Expert VGAT-ChR2 mice. An ambient blue masking light was used in the training sessions as well as testing days. Testing sessions started when mice reached Expert levels of performance (d-prime>1). All the areas of interest were examined in each mouse by inactivating one area per session. The order for the areas was randomized across mice, but inactivations of deep areas (mPFC and dCA1) were performed last. Three sessions per superficial area were performed in each mouse, followed by one session for each deep area. An optic fiber (400 µm; NA = 0.39, Thorlabs) coupled to a 470 nm high power LED (M470F3, Thorlabs, USA) was positioned in contact to the thinned bone for superficial areas or inserted above the left dCA1 at a depth of 1000 µm below the pia. In a subset of mice, dCA1 inactivation was performed using an optrode (silicon probe with an attached optical fiber: 100 µm; NA = 0.22, A1×32-Poly3-10mm-50 s-177-OA32, NeuroNexus, MI, USA). A similar optrode was used for all mPFC inactivations by inserting the tip of the fiber at a depth of 1700 µm, just above the prelimbic area of mPFC. The optrodes were connected to a blue Laser (MBL-F-473/200mW, GMP SA, Switzerland).

Light trials were randomly interleaved with light-off control trials and made up 1/3 of Go and No-Go trials. On light trials, a 100 Hz (40 Hz with laser) train of blue light pulses (50-65% duty cycle, mean power 8-10 mW) was applied in one of the 4 possible windows: *Baseline* (from visual cue onset to 800 ms after), *Whisker* (from 100 ms before the whisker onset to 100 ms after), *Delay* (from 200 ms after the whisker onset to 900 ms after) and *Response* (from auditory cue onset to 1000 ms after). All light windows were terminated by an additional 100 ms ramping down to prevent rebound excitation. In total, 21,293 light trials were tested in 9 mice, 11 areas and 4 trial epochs. On average, for each area and trial epoch, 60.7 ± 6.6 (mean ± SD) light trials were delivered for superficial areas in each mouse across 3 sessions; for deep areas (i.e. mPFC and dCA1), 22.4 ± 4 light trials were examined in one session.

### Histology and localization of electrode/optical fiber tracks

At the end of experiments mice were perfused with phosphate buffered saline (PBS) followed by 4% paraformaldehyde (PFA, Electron Microscopy Science, USA) in PBS. The brain was post-fixed overnight at room temperature. Expression of RCaMP was observed by epifluorescence microscope in serial 100-µm coronal sections cut by a conventional vibratome (VT 1000S; Leica, Wetzlar, Germany). The DiI track of silicon probes were identified with either two-photon tomography (Mayrhofer et al., 2019) or conventional histological analysis. For three-dimensional imaging with two photon tomography, we embedded the brains in 3-5 % oxidized agarose (Type-I agarose, Merck KGaA, Germany) and covalently cross-linked the brain to the agarose by incubating overnight at 4 °C in 0.5 – 1 % sodium borohydride (NaBH_4_, Merck KGaA, Germany) in 0.05 M sodium borate buffer. We imaged the brains in a custom-made two-photon serial microscope, which was controlled using Matlab-based software (ScanImage 2017b, Vidrio Technologies, USA) and BakingTray https://github.com/BaselLaserMouse/BakingTray, version master: 2019/05/20, extension for serial sectioning) (Han et al., 2018). The setup consists of a two-photon microscope coupled with a vibratome (VT1000S, Leica, Germany) and a high-precision X/Y/Z stage (X/Y: V-580; Z: L-310, Physik Instrumente, Germany). The thickness of a physical slice was set to be 50 µm for the entire brain and we acquired optical sections at 25 µm using a high-precision piezo objective scanner (PIFOC P-725, Physik Instrumente, Germany) in two channels (green channel: 500 – 550 nm, ET525/50, Chroma, USA; red channel: 580 – 630 nm, ET605/70, Chroma, USA). Each section was imaged by 7 % overlapping 1025×1025-µm tiles. A 16x water immersion objective lens (LWD 16x/0.80W; MRP07220, Nikon, Japan), with a resolution of 1 µm in X and Y and measured axial point spread function of ∼5 µm full width at half maximum. After image acquisition, the raw images were stitched using a Matlab-based software (StitchIt, https://github.com/BaselLaserMouse/StitchIt). The stitched images were then down-sampled by a factor of 25 in X and Y obtaining a voxel size of 25 × 25 × 25 µm, using a Matlab-based software (MaSIV, https://github.com/alexanderbrown/masiv) to match the Allen Mouse Common Coordinate Framework version 3 (Wang et al., 2020). We used a Matlab-based software (*ARA tools*, https://github.com/SainsburyWellcomeCentre/ara_tools) (Han et al., 2018) to register brain volumes and probe locations to the Allen mouse brain atlas. For some brains with DiI tracks, 100 µm-thick serial sections were cut on a conventional vibratome. The slices were then mounted and imaged under a fluorescence microscope (Leica DM5500). Matlab-based software (*Allen CCF tools*, https://github.com/cortex-lab/allenCCF) was used to register brain slices and probe locations to Allen mouse brain atlas (Shamash et al., 2018).

## QUANTIFICATION AND STATISTICAL ANALYSIS

### Wide-field imaging data

Sessions in which the difference between the Hit rate and False-alarm rate was larger than 0.1 (for Novice) and smaller than 0.2 (for Expert) were excluded from further analysis. In total, 62 Novice sessions and 82 Expert sessions from 7 RCaMP mice, and 57 Expert sessions from 7 tdTomato mice were used for analysis. Acquired images were down-sampled to 77×96 pixels (111 μm/pixel). For each trial, we calculated the normalized signal intensity of each pixel as ΔF/F_0_ = (F-F_0_)/F_0_, where F is the intensity of a pixel in each frame, and F_0_ is the mean intensity of that pixel during the 1 second baseline period before the onset of the visual cue. In each imaging session, mean ΔF/F_0_ images for different trial outcomes (Hit, Miss, False-alarm and Correct-rejection trials) were calculated by averaging all trials in each trial type, or by averaging “Quiet” trials in which mean jaw speed during the 1-second delay period after the whisker stimulus did not exceed 4 times of the mean absolute deviation of the jaw speed (angle) during the 1-second baseline period in each trial. Images from different mice were aligned according to the functionally-identified C2-barrel (RCaMP mice) (Mayrhofer et al., 2019) and the cerebellar tentorium (RCaMP and tdTomato mice), and smoothed by spatial gaussian filter (sigma = 1 pixel, 111 μm). Those trial-averaged images in each session were used as individual samples for statistical analysis. To test statistical differences in the pixel values, Wilcoxon rank-sum test (Expert vs Novice and RCaMP vs tdTomato) or Wilcoxon signed-rank test (Hit vs Miss and Miss vs Correct-rejection) was performed in each pixel, and *p*-value was corrected for multiple comparison by false-discovery rate, FDR (Benjamini and Hochberg, 1995). The corrected *p*-values were log-scaled (-log_10_P) to create spatial *p*-value maps. Borders between anatomical areas were drawn on the functional images (Vanni et al., 2017) by using Allen Mouse Common Coordinate Framework version 3 (CCF) (Lein et al., 2007; Wang et al., 2020) and ARA tools (Han et al., 2018; MacDowell and Buschman, 2020; Musall et al., 2019; Pinto et al., 2019). First, we defined the three-dimensional location of bregma in 25-μm resolution Allen CCF by considering brain structures in the stereotaxic atlas (Paxinos and Franklin, 2019), and the thickness of skull (325 μm) (Soleimanzad et al., 2017). Second, the atlas was rotated by 24 degrees along the rostro-caudal axis. Third, anatomical borders were projected onto the horizontal plane to make a 24-deg tilted border map. Then, the border map was linearly scaled and horizontally shifted to match the functional images of RCaMP mice according to the C2-barrel, bregma, and the anteromedial end of the left hemisphere.

### Electrophysiology data

Spiking activity on each probe was detected and sorted into different clusters using Klusta, an open source spike sorting software suited for dense multielectrode recordings (Rossant et al., 2016). After an automated clustering step, clusters were manually inspected and refined. Single units were categorized as regular spiking (RSU) or fast-spiking neurons based on the duration of the spike waveform, and, in this study, we specifically focus on the putative excitatory RSUs (spike peak-to-baseline > 0.34 ms, 4415 units in 22 Expert and 1604 units in 8 Novice mice). Activity maps in Figure S4D and S6B were computed by averaging the trial-aligned peristimulus time histograms of all excitatory units recorded on the same probe.

### Assessing Expert/Novice and Hit/Miss differences

Statistical difference between mean firing rates of Expert vs Novice (Figures 3E and 5D) and Hit vs Miss (Figures 5E, S4E and S4F) in each area was identified using non-parametric permutation tests in 50-ms bins and *p*-values were corrected by FDR.

### Receiver Operating Characteristic (ROC) analysis

To quantify the selectivity of ROI calcium traces for Go vs No-Go trials we built ROC curves comparing the distribution of calcium activity in bins of 50 ms including only correct trials (Hit and Correct-rejection). Selectivity index was defined by scaling and shifting the area under the ROC curve (AUC) between -1 and 1:

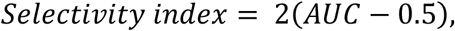

where positive selectivity reflects higher activity in Hits and vice versa (Figure 2D). Similarly, to quantify the selectivity of single units for Go vs No-Go trials we built ROC curves comparing distribution of spiking activity in bins of 100 ms including only correct trials (Hit and Correct-rejection). The area under the ROC curve was then compared to a baseline distribution (5 bins of 100 ms before visual cue onset) to examine the significance of selectivity beyond baseline fluctuations. Non-parametric permutation tests were performed and *p*-values were corrected by FDR and percentage of neurons with significant positive or negative selectivity in each area were identified (p<0.05, FDR-corrected, Figure 3F).

### Clustering neuronal responses

For clustering the neuronal response patterns, RSUs from both Novice and Expert mice (1) with more than 200 spikes throughout the recording, and (2) with more than 5 trials for each trial-type (i.e. Hit, Miss, CR and FA) were included in the analysis (n=5405 out of 6019 RSUs). For each neuron and each trial type, time varying PSTHs (100 ms bin size) were computed over a 4-s window starting from 1 s before the visual cue and lasting until 1 second after the auditory cue. PSTHs from different trial types were baseline subtracted, normalized to the range of values across all bins (of all 4 trial types) and then concatenated resulting in an activity matrix *X* ∈ ℝ^5405 ×160^ whose row *i* corresponds to the concatenated normalized firing rate of the neuron *i* across different trial types (Figure S5A). Other normalization methods such as z-scoring resulted in similar clustering outcomes. To reduce the existing redundancy between firing rate time bins, we used Principal Component Analysis (PCA) and linearly projected firing rate vectors on a low-dimensional space. We applied PCA on the centered version of *X* (i.e. 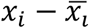.) and found 14 significant components (permutation test with Bonferroni correction for controlling family-wise error rate by 0.05) (Macosko et al., 2015). The weight of different components was equalized by normalizing the data resulting in unity variance for different components (*X*^′^ ∈ ℝ^5405 ×14^).

Next, we employed spectral embedding on the data to detect non-convex and more complex clusters (Abbe, 2017; Von Luxburg, 2007). To do so, we computed the similarity matrix *S* ∈ ℝ^5405 ×5405^ whose element at row *i* and column *j* measures the similarity between *x*_*x*_′ and *x*_*j*_′ as

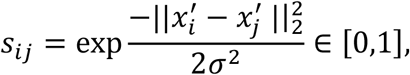

where *σ* is a free parameter determining how local similarity is measured in the feature space. We tuned *σ* by putting the average of similarity values equal to 0.5 (the tuned value for *σ* is 0.0987). Then, we computed the normalized Laplacian matrix as

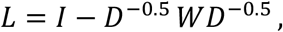

where *I* is the identity matrix, and *D* is the diagonal degree matrix defined as diag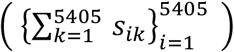. The transformed features are rather abstract and computed as the eigenvectors of *L*. It should be noted that the new feature space is non-linearly transformed version of the PCA-space which is itself a linearly transformed version of the original firing rate space. Such a transformation is believed to naturally separate data points which are clustered together (Abbe, 2017; Von Luxburg, 2007). Using the elbow method on the eigenvalues of matrix *L* (i.e. finding the sharp transition in the derivative of sorted eigenvalues), we considered (after excluding the very 1^st^ eigenvector) the first 13 eigenvectors of matrix *L* as representative features which yielded matrix 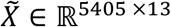.

Finally, neurons were clustered based on the resulting matrix 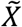 using a Gaussian Mixture Model (GMM). The algorithm considers that underlying distribution of data is a mixture of *K* Gaussians with means {*μ*_1_, …, *μ*_*k*_}, diagonal covariance matrices {Σ _1_, …, Σ _*k*_}, and weights {*p*_1_, …, *p*_*k*_}. For a given *K*, we estimated the parameters of this mixture model by using expected maximization (EM) algorithm (5000 repetitions and 1000 iterations). The number of clusters was then selected (*K* = 24) by minimizing the Bayesian information criterion (BIC) (Engelhard et al., 2019) (Figure S5B). Using the fitted parameters, we assigned a cluster index *c*_*i*_ ∈ {1, …, 24} to each neuron corresponding to the Gaussian distribution to which it belongs with the highest probability. The output of the GMM step was the vector *C* ∈ {1, …, 24}^5405^ containing the cluster indices of neurons. Task-modulated clusters (20/24) were sorted by their onset latency and were labeled based on their task epoch-related response (Figure 3C).

To study to what extent neurons from different brain regions and Novice and Expert mice contribute to the composition of clusters we took 3 steps. First, we quantified the distribution of neurons of each cluster across different brain regions in Novice and Expert mice (Figures 3D, 5F and S5C). To account for the differences in the total number of neurons belonging to each group and brain region, weighted proportions were considered. Next, to identify the patterns which are more prevalent after whisker training, we quantified the percentage of neurons in each cluster that belong to Expert mice (Figure 3D). Similarly, in computing this percentage, weighted proportions were considered to correct for the difference in sample sizes (n=3960 neurons from Expert, n=1445 neurons from Novice). Finally, we defined a “distribution index” which quantifies the spread of each cluster among different brain regions (Figure 3D). For this purpose, we measured the total-variation distance between the weighted distribution of neurons of each cluster across 12 brain regions and the uniform distribution:

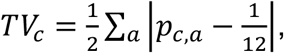

Where *p*_*c,a*_ is the weighted proportion of neurons in cluster *c* belonging to area *a*. Note that *p*_*c,a*_ is normalized with respect to areas, i.e., ∑_*a*_ *p*_*c,a*_ = 1. The distance *TV*_*c*_ takes 0 as its minimum value when the neurons of cluster *c* are uniformly distributed in all areas, and takes 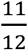 as its maximum value when all neurons of cluster *c* belong to a single brain area. To scale this value between zero and one, for each cluster *c* we defined a distribution index (*D*_*c*_) as:

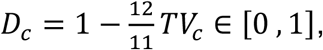

where *D*_*c*_ = 1 indicates that cluster *c* is uniformly distributed among areas, and *D*_*c*_ = 0 indicates that cluster *c* is concentrated in a single brain region.

To characterize changes across learning of the delay task in each area, we computed separately in Novice and Expert mice, the activity pattern of the two most representative clusters (i.e. clusters with the highest number of neurons among all clusters) by averaging the activity among neurons belonging to the pair of area and cluster. The two most representative clusters are labeled as 1st and 2nd rank (Figure S5D).

### Single neuron whisker-evoked response latency

To quantify the latency of whisker-evoked sensory response in spiking activity of single neurons (Figure 5C), we limited the analysis to the first 200-ms window following the whisker stimulus. We first examined whether each neuron was modulated (positively or negatively) in the 200-ms window following the whisker stimulus compared to a 200-ms window prior to the whisker onset. For responsive neurons (*p*<0.05, non-parametric permutation test), latency - calculated on the temporally smoothed PSTHs (1 ms non-overlapping binned PSTH filtered with a Gaussian kernel with *σ*=10 ms) - was defined as the time where the neural activity reached half maximum (half minimum for suppressed neurons) within the 200-ms window. Only responsive neurons are included in the cumulative distributions and boxplots in Figure 5C.

### GLM encoding model

We used Poisson regression to fit an encoding model (generalized linear model, GLM) to predict the spiking activity of each individual neuron given behavioral data (Nelder and Wedderburn, 1972; Park et al., 2014). For each session, we concatenated all correct trials (Hit and Correct-rejection) and then split the data to perform five-fold cross-validation. In Poisson regression, one aims at predicting the spike count y(t) in a time bin t according to the formula:

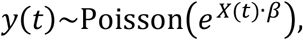

i.e. assuming that the spike counts are sampled from a Poisson distribution with rate that depends on the design matrix *X*(*t*) and on the weight vector *β*. In our case, *y* was constructed by binning the spikes in 100-ms bins. The weights *β* were fit by maximizing the likelihood with Ridge regularization for each fold, and then averaged across the five folds. The parameter that controls the strength of the regularization was determined separately for each neuron using evidence optimization (Cunningham et al., 2008; Park et al., 2014).

The design matrix was constructed by including three types of variables: “event” variables, associated to task-related events; “analog” variables, associated to real-valued behavioral measures from videography; and “slow” variables, which were constant during one trial but could vary over the course of one session. Event variables included the visual cue onset, the whisker stimulus onset, the auditory cue onset and the onset of the first lick. The exact time of lick onset was determined from the high-speed video using a custom algorithm. To assess the delayed effect of such task-related variables, each of these event-like variables was associated with a set of ten 100-ms wide and unit height boxcar basis functions, spanning in total one second after each event. The first-lick variable was associated with two additional boxcar functions covering 0.2 seconds prior to the lick onset, to capture lick-specific preparatory neuronal activity. Analog variables included in the design matrix were the whisker, tongue and jaw speed. These quantities were first extracted from the high-speed videos using custom code and then averaged in 100-ms bins. Among the slow variables, we included the trial index, i.e. a variable that at each trial *k* took a constant value equal to *k*/*k*_*total*_, where *k*_*total*_ is the total number of trials in a session. This variable could capture shifts in a neuron baseline activity due to slow effects across the session such as changes in satiety and motivation. Finally, we included three binary variables that took value one only if the previous trial was an early lick, a False-alarm or a Hit trial, to capture the effect of the previous trial outcome on the subsequent trial. In total, our design matrix had 50 columns, corresponding to the number of free parameters of the model.

To assess the significance of each variable in the design matrix, we fitted a new GLM model obtained by removing the variable of interest (reduced model) from the full model. If for a certain neuron the reduced model fitted the data significantly worse than the full model (*p*<0.05, according to a likelihood ratio test (Buse, 1982)), then that neuron was considered significantly modulated by the removed variable. The reduced model was fitted independently for each fold, using the same data splitting used for the full model. In the likelihood ratio test, the test statistics are given by 2log(*L*_*full*_ / *L*_*reduced*_), where *L*_*full*_ and *L* _*reduced*_ are the full and reduced model likelihood respectively. These statistics were computed for each fold and then averaged to obtain an average statistic, from which the final *p*-value was computed (Buse, 1982). Note that in the presence of correlations among variables, this approach is stringent in that it tends to underestimate the significance of different variables. To separately assess the effect of the onset of event-like variables from their delayed effects, we quantified their significance independently by separately removing the first two basis functions or remaining eight basis functions (Visual, Auditory and Lick). For the whisker variable, since it was very brief in time (10 ms), we removed either the first or the remaining nine bins (referred to as ‘Whisker’ and ‘Delay’ respectively in Figure 6D and 6E). To assess the significance of the modulation due to lick-preparatory neuronal activity we separately removed the two basis functions that preceded the lick onset (referred to as ‘Lick initiation’ in Figure 6D and 6E). Spatial weight maps for selected model variables (Figure S7E) were built by first averaging the weights over the time course of the variable, i.e. by averaging over the weights of the boxcar basis functions. Next, for each neuron these weights were projected on the reconstructed anatomical location in 2D, and were then averaged across all neurons with a certain spatial bin (50×50 µm). The resulting spatial weight map was smoothed using a 2D Gaussian kernel (sigma=150 µm). All the GLM analysis was performed in Matlab using a combination of existing and custom-written code.

### Assessing optogenetic manipulation impact

We measured the impact of optogenetic activation in tjM1 by counting early licks evoked during the delay period. Sessions with a difference between Hit rate and False alarm rate smaller than 0.2 were excluded from the analysis. The early lick rates with the strongest optogenetic stimulation (9 mW) were calculated in each session to test statistical difference between light-off and light trials.

To quantify the impact of optogenetic inactivation we compared mouse averaged performance (n=9; Hit rate, False alarm rate and Early lick rate) for different light windows (i.e. Baseline, Whisker, Delay, Response) to light-off control trials. *P*-values were corrected for multiple comparison (i.e. 4 windows) using Bonferroni correction.

To assess the effect of inactivation on movements, we quantified the change in light versus no-light trials by defining a movement modulation index as:

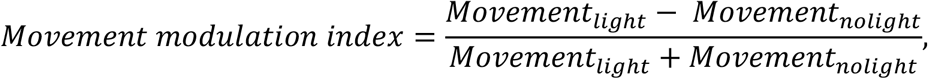

for different orofacial movements (whisker, jaw and tongue speed) and lick spout reading with the piezo sensor (Figure S8).

### Quantifying involvement index

The involvement index was defined by combining the neuronal correlates and behavioral impact of optogenetic inactivation. For each pair of area and temporal window of interest, we built two distributions of bootstrap estimation of the mean, separately for neuronal correlates and inactivation impact, by bootstrapping 1000 times. The neuronal correlates were quantified as the mean firing rate difference in Hit vs Correct-rejection trials across all neurons recorded from 22 Expert mice. The inactivation impact was quantified as the mean change in Hit rate across 9 VGAT-ChR2 mice. The distribution of involvement index was calculated as the product distribution of the two bootstrap distributions.

### Statistics

Data are represented as mean ± SEM unless otherwise noted. The Wilcoxon signed-rank test was used to assess significance in paired comparisons; and the Wilcoxon rank-sum test was used for unpaired comparisons (Matlab implementations). Analysis of spiking activity, selectivity of calcium signals and involvement index was performed using a non-parametric permutation test. The statistical tests used and n numbers are reported explicitly in the main text or figure legends. *P*-values are corrected for multiple comparisons and methods are indicated in figure legends.

**Supplemental Video S1. Wide-field calcium imaging in Hit trials of Novice mice, Related to Figure 2**.

The video shows the cortical activity during Hit trials in Novice RCaMP mice (n=62 sessions). Trial period is indicated in the *upper-right*. The number on the *upper-left* represents the time from visual cue onset, with each frame being 10 ms, and the video shown slowed by a factor of 2.5. ‘+’ depicts bregma. Color scale, -0.5% to +1.5% (ΔF/F0). Scale bar, 1 mm. Whisker stimulus evoked focal activations in the primary and secondary whisker somatosensory cortex, but the cortical activity returned to a baseline level toward the auditory cue.

**Supplemental Video S2. Wide-field calcium imaging in Hit trials of Expert mice, Related to Figure 2**.

The video shows the cortical activity during Hit trials in Expert RCaMP mice (n=82 sessions). Trial period is indicated in the *upper-right*. The number on the *upper-left* represents the time from visual cue onset, with each frame being 10 ms, and the video shown slowed by a factor of 2.5. ‘+’ depicts bregma. Color scale, -0.5% to +1.5% (ΔF/F0). Scale bar, 1 mm. Whisker stimulus evoked activation and deactivation responses across the cortex toward the auditory cue, in clear contrast to Novice mice (Video S1).

**Supplemental Video S3. Wide-field calcium imaging in Miss trials of Novice mice, Related to Figure 2**.

The video shows the cortical activity during Miss trials in Novice RCaMP mice (n=62 sessions). Trial period is indicated in the *upper-right*. The number on the *upper-left* represents the time from visual cue onset, with each frame being 10 ms, and the video shown slowed by a factor of 2.5. ‘+’ depicts bregma. Color scale, -0.5% to +1.5% (ΔF/F0). Scale bar, 1 mm. See also Figure S2. No clear difference in response to the whisker stimulus or during the delay period compared to Hit trials of Novice mice (Video S1).

**Supplemental Video S4. Wide-field calcium imaging in Miss trials of Expert mice, Related to Figure 2**.

The video shows the cortical activity during Miss trials in Expert RCaMP mice (n=82 sessions). Trial period is indicated in the *upper-right*. The number on the *upper-left* represents the time from visual cue onset, with each frame being 10 ms, and the video shown slowed by a factor of 2.5. ‘+’ depicts bregma. Color scale, -0.5% to +1.5% (ΔF/F0). Scale bar, 1 mm. See also Figure S2. Note smaller response amplitudes after the whisker stimulus and during the delay period compared to those in Hit trials of Expert mice (Video S2).

**Video S5. Spiking activity map in Hit trials of Novice mice, Related to Figure 3**.

The movie shows time-lapse maps of mean firing rate in Novice mice in Hit trials. Circles represent different probes in superficial (black outline) and deep (gray outline) areas, and colors show mean z-scored firing rate across the probe at each time window. Trial period is indicated in the *upper-right*. The number on the *upper-left* represents the time from visual cue onset. Each frame is 100 ms, frames are played 5 times slower in time. ‘+’ depicts bregma. Color scale, -0.6 to +0.6 (z-score firing rate) similar to Figure S4D.

**Video S6. Spiking activity map in Hit trials of Expert mice, Related to Figure 3**.

The movie shows time-lapse maps of mean firing rate in Expert mice in Hit trials. Circles represent different probes in superficial (black outline) and deep (gray outline) areas, and colors show mean z-scored firing rate across the probe at each time window. Trial period is indicated in the *upper-right*. The number on the *upper-left* represents the time from visual cue onset. Each frame is 100 ms, frames are played 5 times slower in time. ‘+’ depicts bregma. Color scale, -0.6 to +0.6 (z-score firing rate) similar to Figure S4D. Note higher neuronal activity in frontal regions after the whisker stimulus and during the delay period compared to Novice mice (Video S5).

## Notes

### Competing Interest Statement

The authors have declared no competing interest.

## REFERENCES

Abbe, E. (2017). Community detection and stochastic block models: recent developments. J. Mach. Learn. Res. 18, 6446–6531.

Allen, W.E., Kauvar, I.V., Chen, M.Z., Richman, E.B., Yang, S.J., Chan, K., Gradinaru, V., Deverman, B.E., Luo, L., and Deisseroth, K. (2017). Global representations of goal-directed behavior in distinct cell types of mouse neocortex. Neuron 94, 891–907.

Andermann, M.L., Kerlin, A.M., Roumis, D.K., Glickfeld, L.L., and Reid, R.C. (2011). Functional specialization of mouse higher visual cortical areas. Neuron 72, 1025–1039.

Benjamini, Y., and Hochberg, Y. (1995). Controlling the false discovery rate: a practical and powerful approach to multiple testing. J. R. Stat. Soc. Series B Stat. Methodol. 57, 289–300.

Bethge, P., Carta, S., Lorenzo, D.A., Egolf, L., Goniotaki, D., Madisen, L., Voigt, F.F., Chen, J.L., Schneider, B., Ohkura, M., et al. (2017). An R-CaMP1.07 reporter mouse for cell-type-specific expression of a sensitive red fluorescent calcium indicator. PLoS ONE 12, e0179460.

Buse, A. (1982). The likelihood ratio, Wald, and Lagrange multiplier tests: an expository note. Am. Stat. 36, 153–157.

Buzsáki, G. (2004). Large-scale recording of neuronal ensembles. Nat. Neurosci. 7, 446–451.

Chabrol, F.P., Blot, A., and Mrsic-Flogel, T.D. (2019). Cerebellar contribution to preparatory activity in motor neocortex. Neuron 103, 506–519.

Chen, T.-W., Li, N., Daie, K., and Svoboda, K. (2017). A map of anticipatory activity in mouse motor cortex. Neuron 94, 866–879.

Chikazoe, J., Jimura, K., Hirose, S., Yamashita, K., Miyashita, Y., and Konishi, S. (2009). Preparation to inhibit a response complements response inhibition during performance of a stop-signal task. J. Neurosci. 29, 15870–15877.

Cohen, O., Sherman, E., Zinger, N., Perlmutter, S., and Prut, Y. (2010). Getting ready to move: transmitted information in the corticospinal pathway during preparation for movement. Curr. Opin. Neurobiol. 20, 696–703.

Cover, T.M., and Thomas, J.A. (1991). Entropy, relative entropy and mutual information. In Elements of Information Theory, D. L. Schilling, ed. (John Wiley & Sons, Inc.), pp. 12–49.

Cunningham, J.P., Byron, M.Y., Shenoy, K.V., and Sahani, M. (2008). Inferring neural firing rates from spike trains using Gaussian processes. In Advances in Neural Information Processing Systems 20, P. J. Koller, D. Singer, Y. Roweis, S. Cambridge, eds. (MIT Press), pp. 329–336.

Diamond, M.E., and Arabzadeh, E. (2013). Whisker sensory system – From receptor to decision. Prog. Neurobiol. 103, 28–40.

Duque, J., Greenhouse, I., Labruna, L., and Ivry, R.B. (2017). Physiological markers of motor inhibition during human behavior. Trends Neurosci. 40, 219–236.

Durstewitz, D., Vittoz, N.M., Floresco, S.B., and Seamans, J.K. (2010). Abrupt transitions between prefrontal neural ensemble states accompany behavioral transitions during rule learning. Neuron 66, 438–448.

El-Boustani, S., Sermet, B.S., Foustoukos, G., Oram, T.B., Yizhar, O., and Petersen, C.C.H. (2020). Anatomically and functionally distinct thalamocortical inputs to primary and secondary mouse whisker somatosensory cortices. Nat. Commun. 11, 3342.

Engelhard, B., Finkelstein, J., Cox, J., Fleming, W., Jang, H.J., Ornelas, S., Koay, S.A., Thiberge, S.Y., Daw, N.D., and Tank, D.W. (2019). Specialized coding of sensory, motor and cognitive variables in VTA dopamine neurons. Nature 570, 509–513.

Erlich, J.C., Bialek, M., and Brody, C.D. (2011). A cortical substrate for memory-guided orienting in the rat. Neuron 72, 330–343.

Esmaeili, V., and Diamond, M.E. (2019). Neuronal correlates of tactile working memory in prefrontal and vibrissal somatosensory cortex. Cell Rep. 27, 3167–3181.

Esmaeili, V., Tamura, K., Foustoukos, G., Oryshchuk, A., Crochet, S., and Petersen, C.C.H. (2020). Cortical circuits for transforming whisker sensation into goal-directed licking. Curr. Opin. Neurobiol. 65, 38–48.

Fassihi, A., Akrami, A., Pulecchi, F., Schönfelder, V., and Diamond, M.E. (2017). Transformation of perception from sensory to motor cortex. Curr. Biol. 27, 1585–1596.

Ferezou, I., Haiss, F., Gentet, L.J., Aronoff, R., Weber, B., and Petersen, C.C.H. (2007). Spatiotemporal dynamics of cortical sensorimotor integration in behaving mice. Neuron 56, 907–923.

Funahashi, S., Bruce, C.J., and Goldman-Rakic, P.S. (1989). Mnemonic coding of visual space in the monkey’s dorsolateral prefrontal cortex. J. Neurophysiol. 61, 331– 349.

Fuster, J.M., and Alexander, G.E. (1971). Neuron activity related to short-term memory. Science 173, 652–654.

Gao, Z., Davis, C., Thomas, A.M., Economo, M.N., Abrego, A.M., Svoboda, K., De Zeeuw, C.I., and Li, N. (2018). A cortico-cerebellar loop for motor planning. Nature 563, 113–116.

Gerstner, W., Kistler, W.M., Naud, R., and Paninski, L. (2014). Neuronal dynamics: from single neurons to networks and models of cognition (Cambridge University Press).

Gilad, A., Gallero-Salas, Y., Groos, D., and Helmchen, F. (2018). Behavioral strategy determines frontal or posterior location of short-term memory in neocortex. Neuron 99, 814–828.

Gong, S., Doughty, M., Harbaugh, C.R., Cummins, A., Hatten, M.E., Heintz, N., and Gerfen, C.R. (2007). Targeting Cre recombinase to specific neuron populations with bacterial artificial chromosome constructs. J. Neurosci. 27, 9817–9823.

Gorski, J.A., Talley, T., Qiu, M., Puelles, L., Rubenstein, J.L.R., and Jones, K.R. (2002). Cortical excitatory neurons and glia, but not GABAergic neurons, are produced in the Emx1-expressing lineage. J. Neurosci. 22, 6309–6314.

Guo, Z.V., Li, N., Huber, D., Ophir, E., Gutnisky, D., Ting, J.T., Feng, G., and Svoboda, K. (2014). Flow of cortical activity underlying a tactile decision in mice. Neuron 81, 179– 194.

Guo, Z.V., Inagaki, H.K., Daie, K., Druckmann, S., Gerfen, C.R., and Svoboda, K. (2017). Maintenance of persistent activity in a frontal thalamocortical loop. Nature 545, 181–186.

Han, Y., Kebschull, J.M., Campbell, R.A.A., Cowan, D., Imhof, F., Zador, A.M., and Mrsic-Flogel, T.D. (2018). The logic of single-cell projections from visual cortex. Nature 556, 51–56.

Harvey, C.D., Coen, P., and Tank, D.W. (2012). Choice-specific sequences in parietal cortex during a virtual-navigation decision task. Nature 484, 62–68.

Hastie, T., Tibshirani, R., and Friedman, J. (2009). The elements of statistical learning: data mining, inference, and prediction (Springer Science & Business Media).

Hattori, R., Danskin, B., Babic, Z., Mlynaryk, N., and Komiyama, T. (2019). Area-specificity and plasticity of history-dependent value coding during learning. Cell 177, 1858–1872.

Komiyama, T., Sato, T.R., O’Connor, D.H., Zhang, Y.-X., Huber, D., Hooks, B.M., Gabitto, M., and Svoboda, K. (2010). Learning-related fine-scale specificity imaged in motor cortex circuits of behaving mice. Nature 464, 1182–1186.

Kwon, S.E., Yang, H., Minamisawa, G., and O’Connor, D.H. (2016). Sensory and decision-related activity propagate in a cortical feedback loop during touch perception. Nat. Neurosci. 19, 1243–1249.

Kyriakatos, A., Sadashivaiah, V., Zhang, Y., Motta, A., Auffret, M., and Petersen, C.C.H. (2017). Voltage-sensitive dye imaging of mouse neocortex during a whisker detection task. Neurophotonics 4, 031204.

de Lafuente, V., and Romo, R. (2006). Neural correlate of subjective sensory experience gradually builds up across cortical areas. Proc. Natl. Acad. Sci. USA 103, 14266–14271.

Le Merre, P., Esmaeili, V., Charrière, E., Galan, K., Salin, P.-A., Petersen, C.C.H., and Crochet, S. (2018). Reward-based learning drives rapid sensory signals in medial prefrontal cortex and dorsal hippocampus necessary for goal-directed behavior. Neuron 97, 83–91.

Lefort, S., Tomm, C., Floyd Sarria, J.-C., and Petersen, C.C.H. (2009). The excitatory neuronal network of the C2 barrel column in mouse primary somatosensory cortex. Neuron 61, 301–316.

Lein, E.S., Hawrylycz, M.J., Ao, N., Ayres, M., Bensinger, A., Bernard, A., Boe, A.F., Boguski, M.S., Brockway, K.S., Byrnes, E.J., et al. (2007). Genome-wide atlas of gene expression in the adult mouse brain. Nature 445, 168–176.

Li, N., Chen, T.-W., Guo, Z.V., Gerfen, C.R., and Svoboda, K. (2015). A motor cortex circuit for motor planning and movement. Nature 519, 51–56.

Luo, P., Li, A., Zheng, Y., Han, Y., Tian, J., Xu, Z., Gong, H., and Li, X. (2019). Whole brain mapping of long-range direct input to glutamatergic and GABAergic neurons in motor cortex. Front. Neuroanat. 13, 44.

MacDowell, C.J., and Buschman, T.J. (2020). Low-dimensional spatiotemporal dynamics underlie cortex-wide neural activity. Curr. Biol. 30, 2665–2680.

Macosko, E.Z., Basu, A., Satija, R., Nemesh, J., Shekhar, K., Goldman, M., Tirosh, I., Bialas, A.R., Kamitaki, N., and Martersteck, E.M. (2015). Highly parallel genome-wide expression profiling of individual cells using nanoliter droplets. Cell 161, 1202–1214.

Madisen, L., Zwingman, T.A., Sunkin, S.M., Oh, S.W., Zariwala, H.A., Gu, H., Ng, L.L., Palmiter, R.D., Hawrylycz, M.J., Jones, A.R., et al. (2010). A robust and high-throughput Cre reporting and characterization system for the whole mouse brain. Nat. Neurosci. 13, 133–140.

Madisen, L., Mao, T., Koch, H., Zhuo, J., Berenyi, A., Fujisawa, S., Hsu, Y.-W.A., Garcia, A.J., Gu, X., Zanella, S., et al. (2012). A toolbox of Cre-dependent optogenetic transgenic mice for light-induced activation and silencing. Nat. Neurosci. 15, 793–802.

Makino, H., Ren, C., Liu, H., Kim, A.N., Kondapaneni, N., Liu, X., Kuzum, D., and Komiyama, T. (2017). Transformation of cortex-wide emergent properties during motor learning. Neuron 94, 880–890.

Mao, T., Kusefoglu, D., Hooks, B.M., Huber, D., Petreanu, L., and Svoboda, K. (2011). Long-range neuronal circuits underlying the interaction between sensory and motor cortex. Neuron 72, 111–123.

Marshel, J.H., Garrett, M.E., Nauhaus, I., and Callaway, E.M. (2011). Functional specialization of seven mouse visual cortical areas. Neuron 72, 1040–1054.

Mayford, M., Bach, M.E., Huang, Y.Y., Wang, L., Hawkins, R.D., and Kandel, E.R. (1996). Control of memory formation through regulated expression of a CaMKII transgene. Science 274, 1678–1683.

Mayrhofer, J.M., El-Boustani, S., Foustoukos, G., Auffret, M., Tamura, K., and Petersen, C.C.H. (2019). Distinct contributions of whisker sensory cortex and tongue-jaw motor cortex in a goal-directed sensorimotor transformation. Neuron 103, 1034– 1043.

Miyashita, T., and Feldman, D.E. (2013). Behavioral detection of passive whisker stimuli requires somatosensory cortex. Cereb. Cortex 23, 1655–1662.

Musall, S., Kaufman, M.T., Juavinett, A.L., Gluf, S., and Churchland, A.K. (2019). Single-trial neural dynamics are dominated by richly varied movements. Nat. Neurosci. 22, 1677–1686.

Nelder, J.A., and Wedderburn, R.W.M. (1972). Generalized linear models. J. R. Stat. Soc. Series A Gen. 135, 370–384.

Oh, S.W., Harris, J.A., Ng, L., Winslow, B., Cain, N., Mihalas, S., Wang, Q., Lau, C., Kuan, L., Henry, A.M., et al. (2014). A mesoscale connectome of the mouse brain. Nature 508, 207–214.

Park, I.M., Meister, M.L.R., Huk, A.C., and Pillow, J.W. (2014). Encoding and decoding in parietal cortex during sensorimotor decision-making. Nat. Neurosci. 17, 1395–1403.

Paxinos, G., and Franklin, K.B. (2019). Paxinos and Franklin’s the mouse brain in stereotaxic coordinates (Academic press).

Petersen, C.C.H. (2019). Sensorimotor processing in the rodent barrel cortex. Nat. Rev. Neurosci. 20, 533–546.

Pinto, L., Rajan, K., DePasquale, B., Thiberge, S.Y., Tank, D.W., and Brody, C.D. (2019). Task-dependent changes in the large-scale dynamics and necessity of cortical regions. Neuron 104, 810–824.

Popescu, A.T., Zhou, M.R., and Poo, M.-M. (2016). Phasic dopamine release in the medial prefrontal cortex enhances stimulus discrimination. Proc. Natl. Acad. Sci. USA 113, E3169–3176.

Powell, N.J., and Redish, A.D. (2016). Representational changes of latent strategies in rat medial prefrontal cortex precede changes in behaviour. Nat. Commun. 7, 12830.

Rossant, C., Kadir, S.N., Goodman, D.F.M., Schulman, J., Hunter, M.L.D., Saleem, A.B., Grosmark, A., Belluscio, M., Denfield, G.H., Ecker, A.S., et al. (2016). Spike sorting for large, dense electrode arrays. Nat. Neurosci. 19, 634–641.

Sachidhanandam, S., Sreenivasan, V., Kyriakatos, A., Kremer, Y., and Petersen, C.C.H. (2013). Membrane potential correlates of sensory perception in mouse barrel cortex. Nat. Neurosci. 16, 1671–1677.

Shamash, P., Carandini, M., Harris, K., and Steinmetz, N. (2018). A tool for analyzing electrode tracks from slice histology. bioRxiv 447995.

Sippy, T., Lapray, D., Crochet, S., and Petersen, C.C.H. (2015). Cell-type-specific sensorimotor processing in striatal projection neurons during goal-directed behavior. Neuron 88, 298–305.

Soleimanzad, H., Gurden, H., and Pain, F. (2017). Optical properties of mice skull bone in the 455-to 705-nm range. J. Biomedi. Opt. 22, 10503.

Sreenivasan, V., Esmaeili, V., Kiritani, T., Galan, K., Crochet, S., and Petersen, C.C.H. (2016). Movement initiation signals in mouse whisker motor cortex. Neuron 92, 1368– 1382.

Sreenivasan, V., Kyriakatos, A., Mateo, C., Jaeger, D., and Petersen, C.C.H. (2017). Parallel pathways from whisker and visual sensory cortices to distinct frontal regions of mouse neocortex. Neurophotonics 4, 031203.

Steinmetz, N.A., Zatka-Haas, P., Carandini, M., and Harris, K.D. (2019). Distributed coding of choice, action and engagement across the mouse brain. Nature 576, 266– 273.

Stringer, C., Pachitariu, M., Steinmetz, N., Reddy, C.B., Carandini, M., and Harris, K.D. (2019). Spontaneous behaviors drive multidimensional, brainwide activity. Science 364, 255.

Taniguchi, H., He, M., Wu, P., Kim, S., Paik, R., Sugino, K., Kvitsiani, D., Kvitsani, D., Fu, Y., Lu, J., et al. (2011). A resource of Cre driver lines for genetic targeting of GABAergic neurons in cerebral cortex. Neuron 71, 995–1013.

Tanji, J., and Evarts, E.V. (1976). Anticipatory activity of motor cortex neurons in relation to direction of an intended movement. J. Neurophysiol. 39, 1062–1068.

Vanni, M.P., Chan, A.W., Balbi, M., Silasi, G., and Murphy, T.H. (2017). Mesoscale mapping of mouse cortex reveals frequency-dependent cycling between distinct macroscale functional modules. J. Neurosci. 37, 7513–7533.

Von Luxburg, U. (2007). A tutorial on spectral clustering. Stat. Comput. 17, 395–416.

Wang, Q., and Burkhalter, A. (2007). Area map of mouse visual cortex. J. Comp. Neurol. 502, 339–357.

Wang, Q., Ding, S.-L., Li, Y., Royall, J., Feng, D., Lesnar, P., Graddis, N., Naeemi, M., Facer, B., Ho, A., et al. (2020). The Allen mouse brain common coordinate framework: a 3D reference atlas. Cell 181, 936–953.

Yamashita, T., and Petersen, C.C.H. (2016). Target-specific membrane potential dynamics of neocortical projection neurons during goal-directed behavior. eLife 5, e15798.

Yang, H., Kwon, S.E., Severson, K.S., and O’Connor, D.H. (2016). Origins of choice-related activity in mouse somatosensory cortex. Nat. Neurosci. 19, 127–134.

Zhao, S., Ting, J.T., Atallah, H.E., Qiu, L., Tan, J., Gloss, B., Augustine, G.J., Deisseroth, K., Luo, M., Graybiel, A.M., et al. (2011). Cell type–specific channelrhodopsin-2 transgenic mice for optogenetic dissection of neural circuitry function. Nat. Methods 8, 745–752.

