## Supplemental Figures and Table for "Rapid suppression and sustained activation of distinct cortical regions for a delayed sensory-triggered motor response"

Supplemental Information consists of:

Supplemental Figure S1, related to Figure 1

Supplemental Figure S2, related to Figure 2

Supplemental Figure S3, related to Figure 2

Supplemental Figure S4, related to Figure 3

Supplemental Figure S5, related to Figures 3 and 5

Supplemental Figure S6, related to Figure 5

Supplemental Figure S7, related to Figure 6

Supplemental Figure S8, related to Figure 7

Supplemental Table S1, related to Figure 6

### Supplemental Figure S1

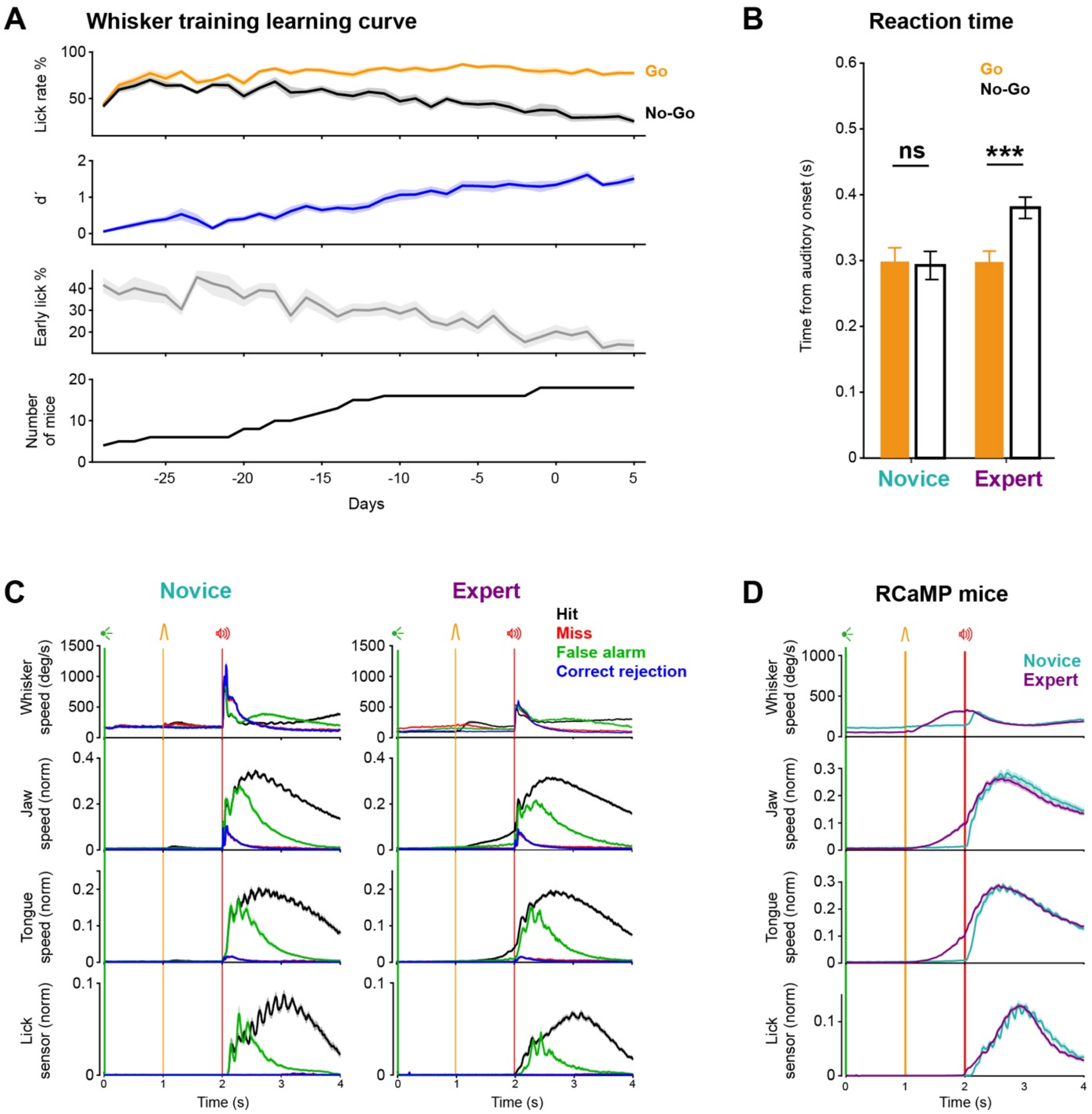

**Figure S1. Whisker training changes behavioral patterns, Related to Figure 1.**

(A) Time courses of behavioral performance across whisker training. From *top* to *bottom*: Lick rate (i.e. the probability of licking in the Response window) in Go and No-Go trials, discriminability of Go and No-Go trials (d-prime), percentage of early licks (licks between visual and auditory cues), and number of mice in each day are plotted along training days relative to the start of neuronal recording (day=0). Only mice used for electrophysiology are shown here (n=18 mice).

(B) Changes in licking behavior. Reaction time (first lick time relative to auditory cue onset time) in completed trials. Reaction times were shorter in Go (Hit, orange) compared to No-Go (False-alarm, black) trials in Expert ($p<0.01$, n=25; Wilcoxon signed-rank test), but not in Novice mice ($p=0.14$, n=15). Mice used for electrophysiology and imaging are included.

(C) Orofacial movements in different trial types. Average movement across mice (mean ± SEM). Novice mice (n=8) showed similar levels of movements across different trial types until auditory cue whereas Expert mice (n=18) increased movements of tongue and jaw toward the auditory cue selectively in Hit trials. Only mice used for electrophysiology are included.

(D) Orofacial movements in Hit trials for Novice and Expert RCaMP mice (n=7). Same configuration as Figure 1F, *left*.

**Supplemental Figure S2**

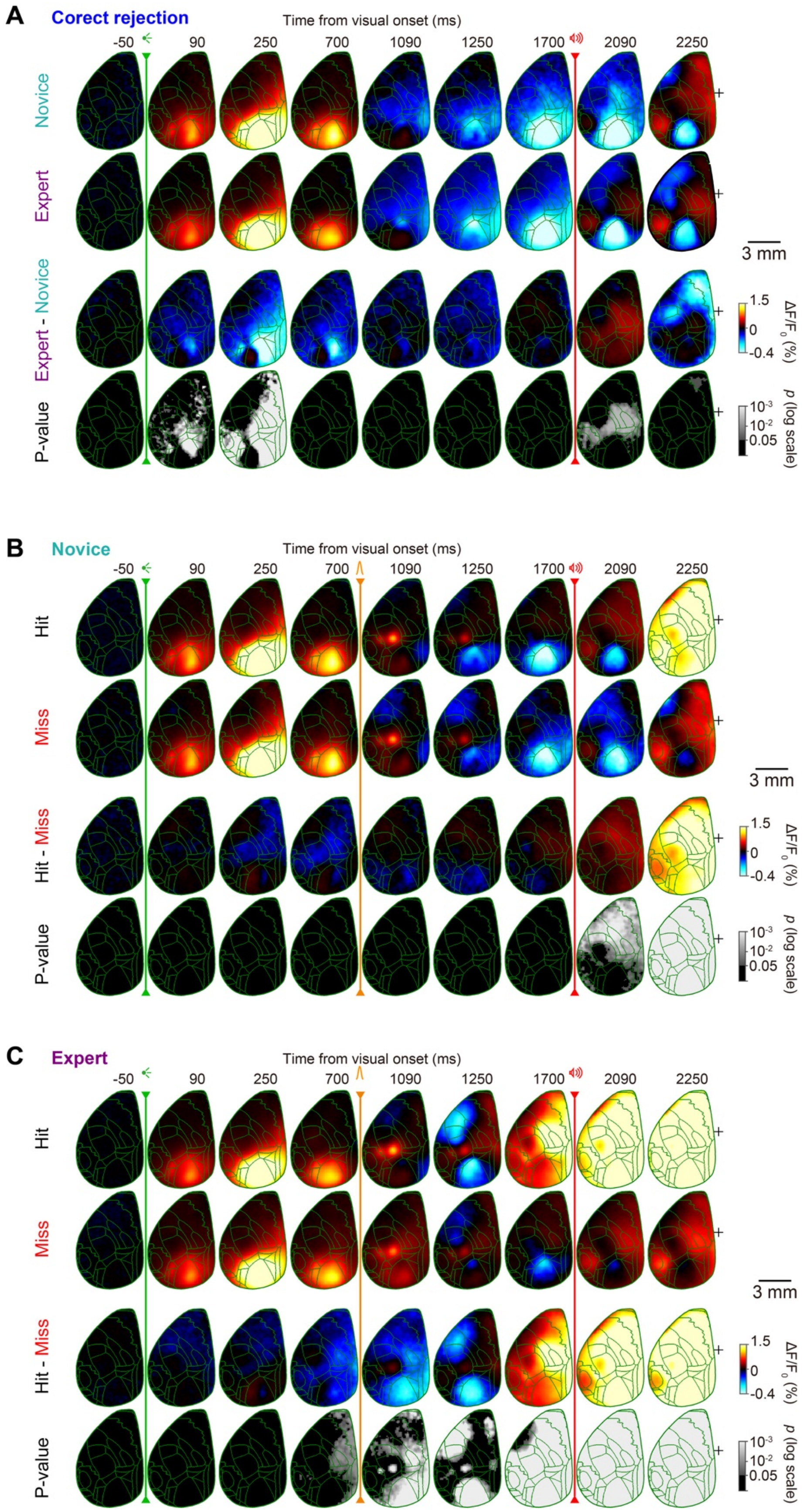

**Figure S2. Wide-field imaging in different trial types, Related to Figure 2.**

(A) Time-course of global cortical activity in Correct-rejection trials for Novice vs Expert mice. Each frame shows instantaneous $\Delta F/F_0$ without averaging (10 ms/frame). For each pixel, baseline activity in a 50 ms window before visual cue onset was subtracted. Mean calcium activity of 62 Novice and 82 Expert sessions from 7 mice, Novice and Expert difference, and the statistical significance of the difference (*p*-value of Wilcoxon rank-sum test, FDR-corrected) are plotted from *top* to *bottom*. Green traces, anatomical borders based on Allen Mouse Brain Atlas. Black '+' indicates bregma.

(B-C) Time-course of global cortical activity in Novice (B) and Expert (C) mice comparing Hit vs Miss trials. Same configuration as panel A. Mean calcium activity of 62 Novice and 82 Expert sessions from 7 mice. Hit, Miss, Hit-Miss difference, and the statistical significance of the difference (*p*-value of Wilcoxon signed-rank test, FDR-corrected) are plotted from *top* to *bottom*.

### Supplemental Figure S3

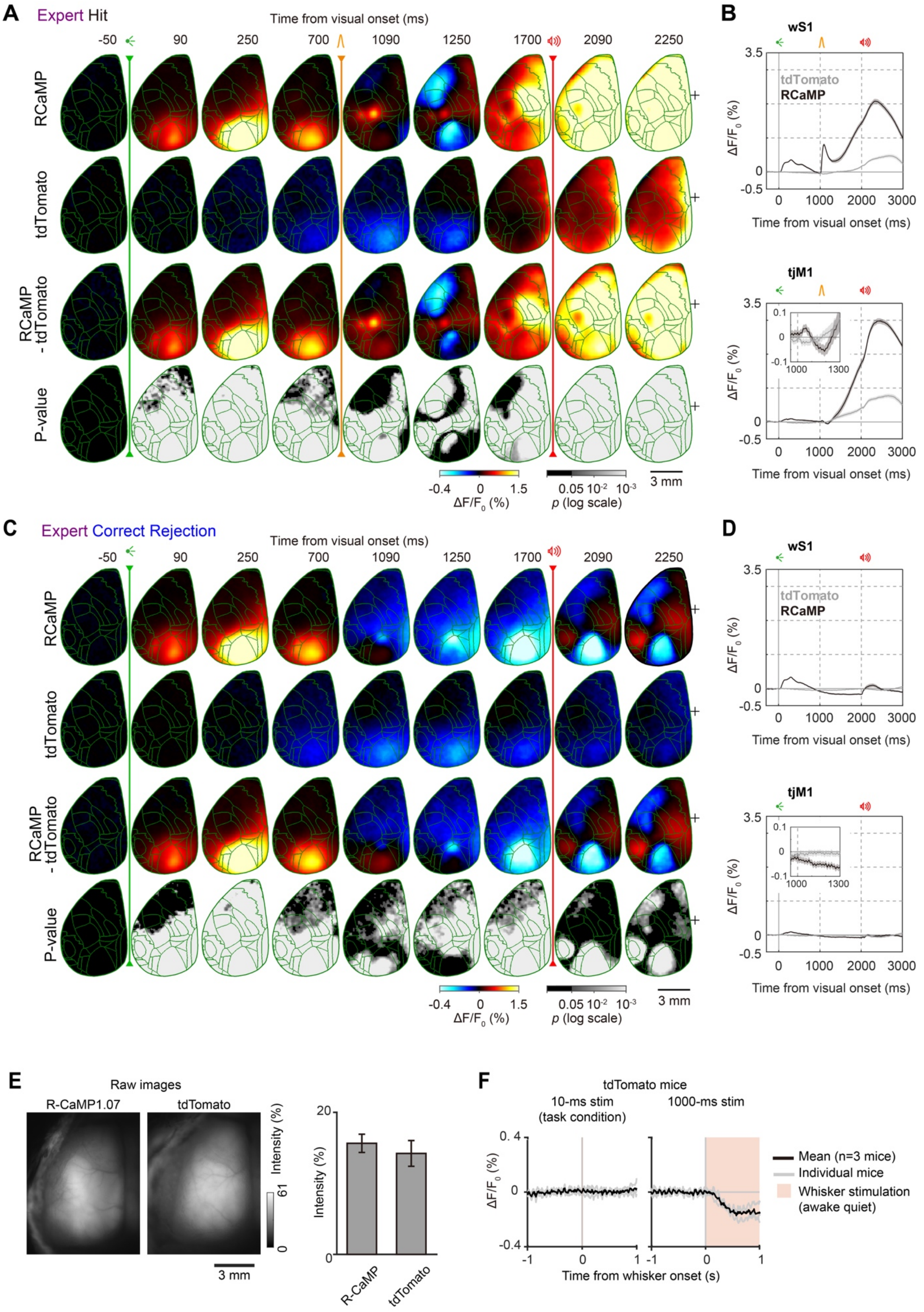

**Figure S3. Control experiment in tdTomato mice, Related to Figure 2.**

(A-D) Time-course of cortical fluorescence in control mice expressing a red fluorescence protein tdTomato during Hit (A-B) and Correct-rejection (C-D) trials, compared with RCaMP mice.

A and C, each frame shows instantaneous $\Delta F/F_0$ without averaging (10 ms/frame). For each pixel, baseline activity in a 50 ms window before visual cue onset was subtracted. RCaMP (mean of 82 Expert sessions from 7 mice) and tdTomato (mean of 57 sessions from 7 Expert mice), RCaMP-tdTomato difference, and the statistical significance of the difference (*p*-value of Wilcoxon rank-sum test, FDR-corrected) are plotted from *top* to *bottom*. Green traces, anatomical borders based on Allen Mouse Brain Atlas. Black '+' indicates bregma. Note slow and spatially diffuse signals related to hemodynamics in tdTomato mice. After subtraction of the tdTomato data, RCaMP mice still show similar patterns of cortical responses.

B and D, fluorescence traces (mean ± SEM) in wS1 (*top*) and tjM1 (*bottom*). ROI size, 7×7 pixels. Inset in tjM1shows enlarged traces right after the whisker onset.

(E) Representative raw fluorescence images of a RCaMP mouse and a tdTomato mouse obtained during wide-field imaging (*left*). There was no statistical difference in the raw intensity between two mouse lines in the imaging conditions for each line (n=7 RCaMP mice and n=7 tdTomato mice, *p*=0.80, Wilcoxon's rank-sum test) (*right*).

(F) Hemodynamic signal in tdTomato mice evoked by whisker stimulation. The brief whisker stimulation (a single 10 ms pulse) used in the task did not evoke detectable changes in wS1, but a prolonged whisker stimulation (100 pulses each of 10 ms at 100 Hz lasting 1000 ms) evoked a strong reduction of cortical fluorescence (n = 3 tdTomato mice).

**Supplemental Figure S4**

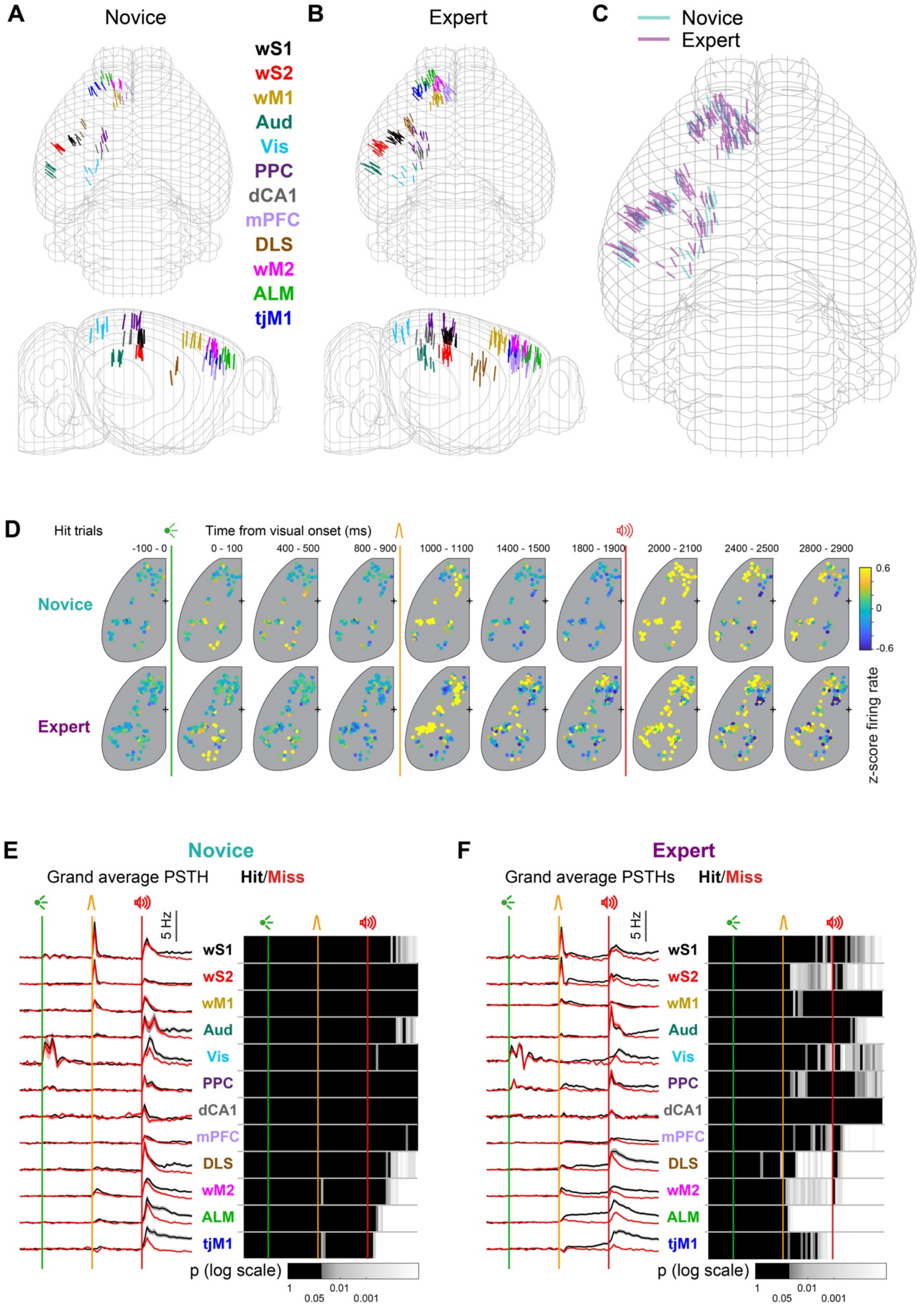

**Figure S4. Silicon probe locations, average activity patterns across probes and firing differences in Hit vs Miss trials in different areas, Related to Figure 3.**

(A-C) Reconstructed location of silicon probes registered to Allen Mouse Brain Atlas in 3D in Novice (A), Expert (B) and overlay of Expert and Novice mice (C).

(D) Time-lapse maps of mean firing rate in Novice and Expert mice in Hit trials. Circles represent different probes and colors show mean z-scored firing rate across the probe at each time window. Neuronal activity patterns are strikingly different between Novice and Expert mice during the delay period (1400 –1900 ms after visual cue onset). Probes from all mice in each group (8 Novice and 22 Expert) are superimposed.

(E) Population firing rate in Novice mice comparing correct vs incorrect Go trials. *Left*, baseline-subtracted (1 s prior to visual cue onset) mean firing rate across cortical areas in Hit and Miss trials of Novice mice are overlaid. *Right*, *p*-value of Hit/Miss comparison in 50 ms consecutive windows (non-parametric permutation test, FDR-corrected) (*right*).

(F) Same as E but for Expert mice. Note prominent differences during the delay period after whisker stimulus across many regions including wS2, DLS, wM2, ALM and tjM1.

### Supplemental Figure S5

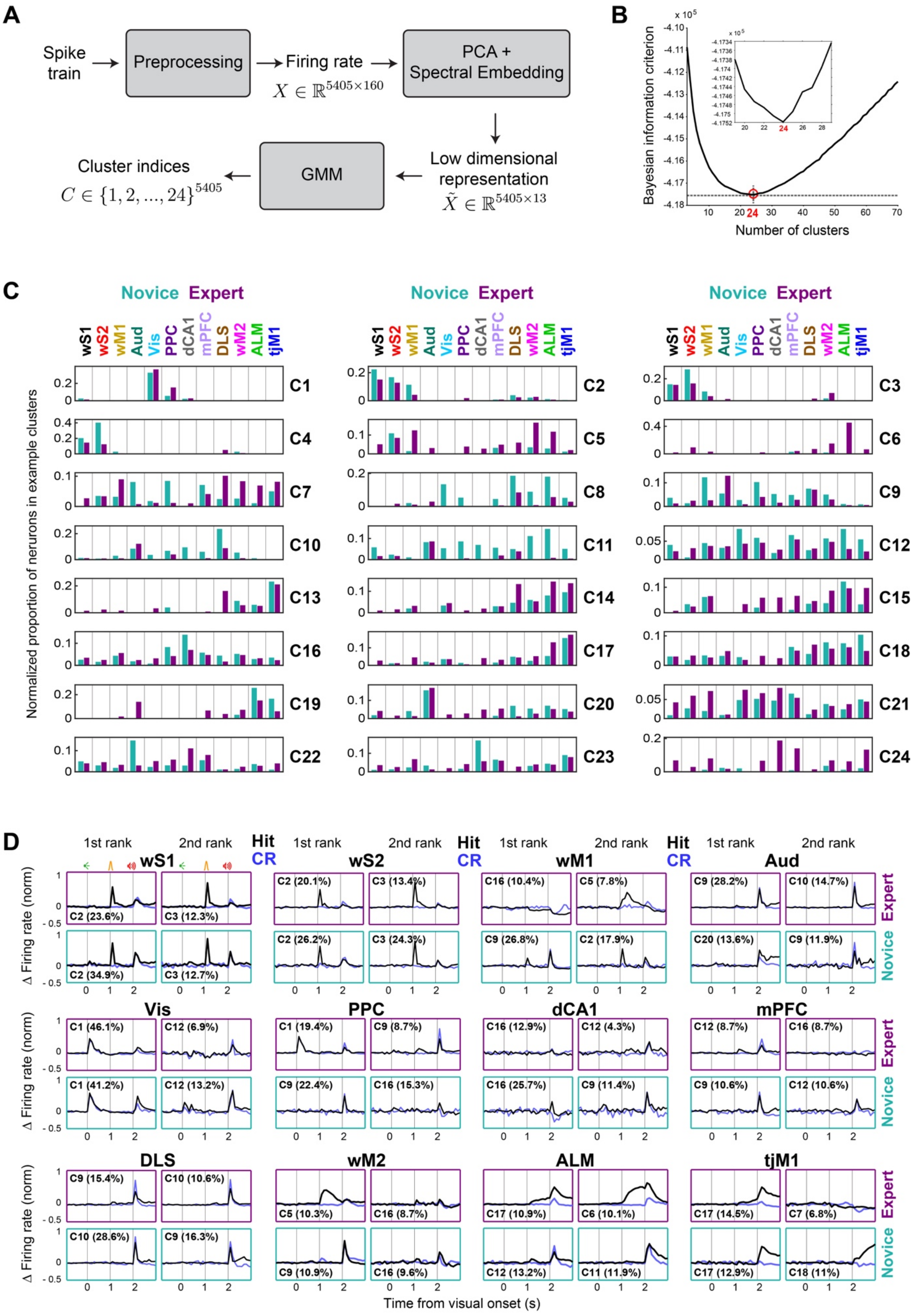

**Figure S5. Unsupervised neuronal clustering, Related to Figures 3 and 5.**

(A) Block diagram indicating the different steps for unsupervised neuronal clustering. Dimensionality reduction and spectral embedding were applied on concatenated trial-type averaged PSTHs of neurons and the results were clustered by fitting a Gaussian mixture model (GMM).

(B) Determination of the number of clusters. Number of optimal clusters (n=24) was determined as the minimum of the Bayesian information criterion (BIC) curve. Inset shows magnified version of BIC values around n=24.

(C) Spatial distribution of clusters across cortex. Weighted proportion of neurons belonging to different cortical regions (similar to rows of heatmaps in Figure 3D but for all 24 clusters). Sorted (from *left* to *right*, and then *top* to *bottom*) based on latency of response onset.

(D) Most prominent firing patterns in different brain regions. For each area the mean firing rate in Hit (black) and Correction rejection (blue) trials are superimposed for the neurons belonging to their two most representative clusters (1st and 2nd rank). Expert (*top*) and Novice (*bottom*) mice. The cluster number is indicated within each frame, as well as the percentage of neurons from the corresponding area belonging to this cluster in parenthesis.

### Supplemental Figure S6

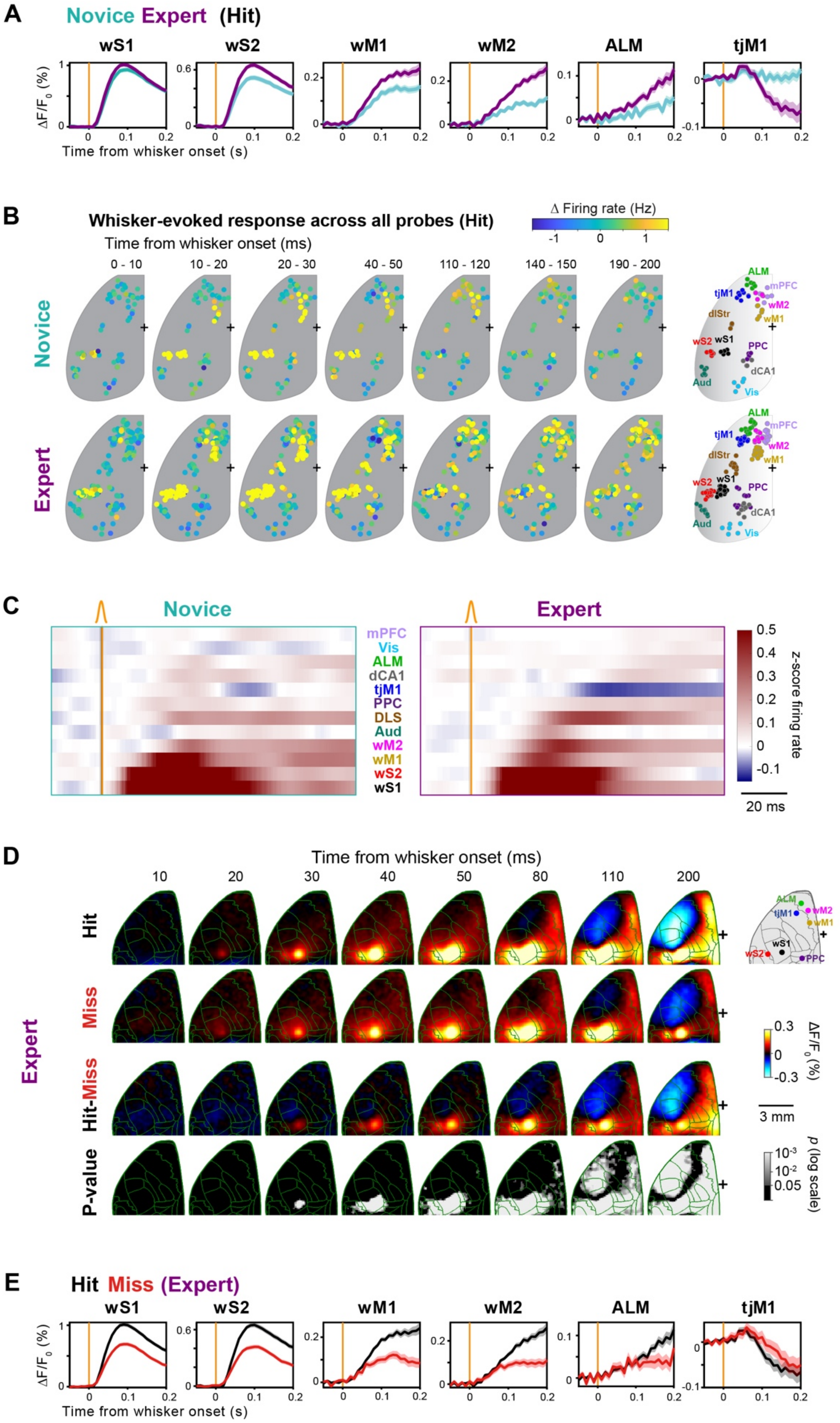

**Figure S6. Critical early delay processing, Related to Figure 5.**

(A) Time courses of whisker-evoked calcium signal in selected regions of interest from the data in Figure 5A. Mean of 62 Novice and 82 Expert sessions from 7 mice (± SEM). ROI size, 3×3 pixels. Activity did not change by learning in the input node wS1, but did diverge in other regions. Note sharp decrease of signal in tjM1 of Expert mice.

(B) Time-lapse maps of mean firing rate immediately after whisker onset. Novice and Expert mice in Hit trials. Same configuration as Figure S4D but with higher temporal resolution (10 ms) for 0-200 ms after whisker onset. Note the propagation of excitation in wS1 → wS2 → wM1 → wM2 → ALM and transient inhibition of tjM1 in Expert mice.

(C) Sequential propagation of whisker-evoked neuronal response in Hit trials (*left*, Novice; *right*, Expert). Mean z-scored firing rate in the first 100 ms window after whisker stimulus are shown. Brain regions are sorted based on their population-average onset latency in Expert mice.

(D) Wide-field signal immediately after whisker stimulus in Hit and Miss trials. Same as Figure 5A but for Hit vs Miss trials of Expert mice (n=82 sessions). From *top* to *bottom*, average calcium signal in Hit, Miss, Hit-Miss difference, and the statistical significance of the comparison (*p*-value of Wilcoxon signed-rank test, FDR-corrected). The schematic (*top-right*) shows the location of regions of interest plotted in E.

(E) Time courses of whisker-evoked calcium signal for Hit and Miss trials in selected regions of interest from the data in D. ROI size, 3×3 pixels.

### Supplemental Figure S7

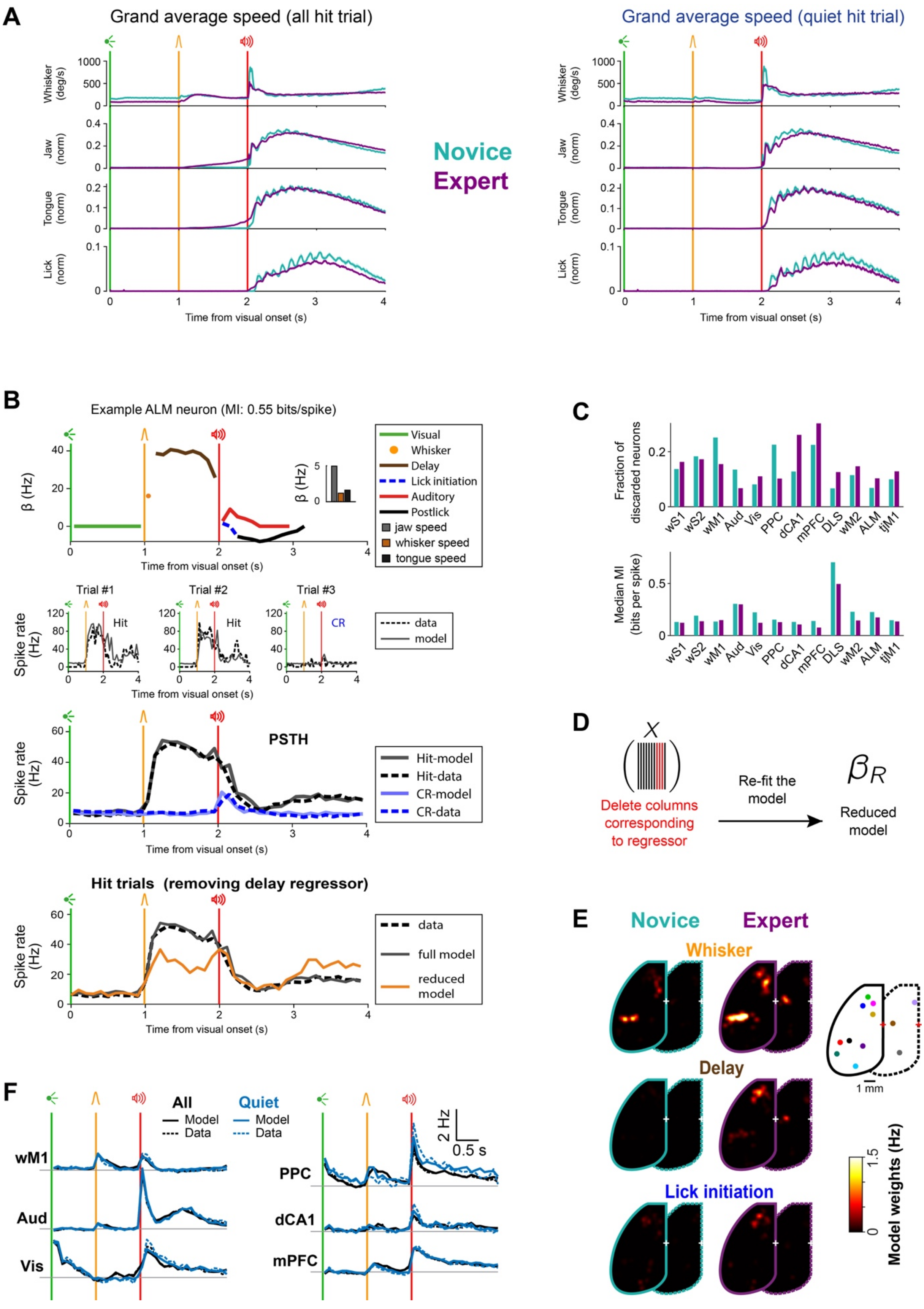

**Figure S7. Orofacial movements in Quiet trials and Poisson encoding model, Related to Figure 6.**

(A) Orofacial movements in selected Quiet trials. The same configuration as Figure 1F, *left*. *Left*, grand average movements in all Hit trials without selection (same data as Figure 1F). *Right*, grand average movements for selected Quiet trials where mice did not show jaw movements. Note that preparatory movements in the delay period after whisker stimulus disappeared.

(B) Model fit for an example ALM neuron. *Top*, model weights ($\beta$) for event variables along the trial timeline are plotted. *Inset*, shows analog variable weights. Weights are shown in units of firing rate (Hz). *Middle*, the overlay of example single-trial firing rate reconstructed from the model (solid lines) and data (dotted lines) are shown for two example Hit (black) and one Correct-rejection (blue) trials. *Bottom*, Average Hit (black) and Correct-rejection (blue) firing rate (PSTHs) reconstructed from the model (plain lines) overlaid with observed data (dotted lines). Orange trace shows PSTH for reduced model after removing delay regressor. Note the high performance of model in reconstructing single trial firing rates and average PSTH.

(C) Model performance. *Top*, fraction of discarded neurons. For each neuron, fit quality was assessed using mutual information (MI) (Cover and Thomas, 1991; Gerstner et al., 2014) (see STAR Methods), a measure of the difference between the fitted model and constant Poisson model capturing only the mean firing rate. Neurons for which the fitted model did not perform better than the constant model (i.e. MI<=0) were excluded from the rest of analysis. Note the higher proportion of excluded neurons in Novice mice, suggesting that neurons became more task-related after whisker training. *Bottom*, median MI values across different regions for Expert and Novice mice.

(D) Evaluating significant contribution of individual predictors. For each fitted neuron, contribution of a model predictor was evaluated by refitting the model after excluding that predictor (reduced model; see panel B, orange PSTH) and comparing it to the full model ($p$<0.05, likelihood ratio test).

(E) Average weight maps of Whisker, Delay and Lick initiation model variables. For each model variable, average model weight map across neurons of superficial (solid border) and deep (dotted border) brain regions of Expert and Novice mice are shown.

(F) Comparison of PSTHs of Quiet trials reconstructed from the GLM model (solid lines) with empirical data (dotted lines). Same configuration as Figure 6F but for the other brain regions. Note that the model fitted to all trials (black) reconstructs well Quiet (blue) trials.

### Supplemental Figure S8

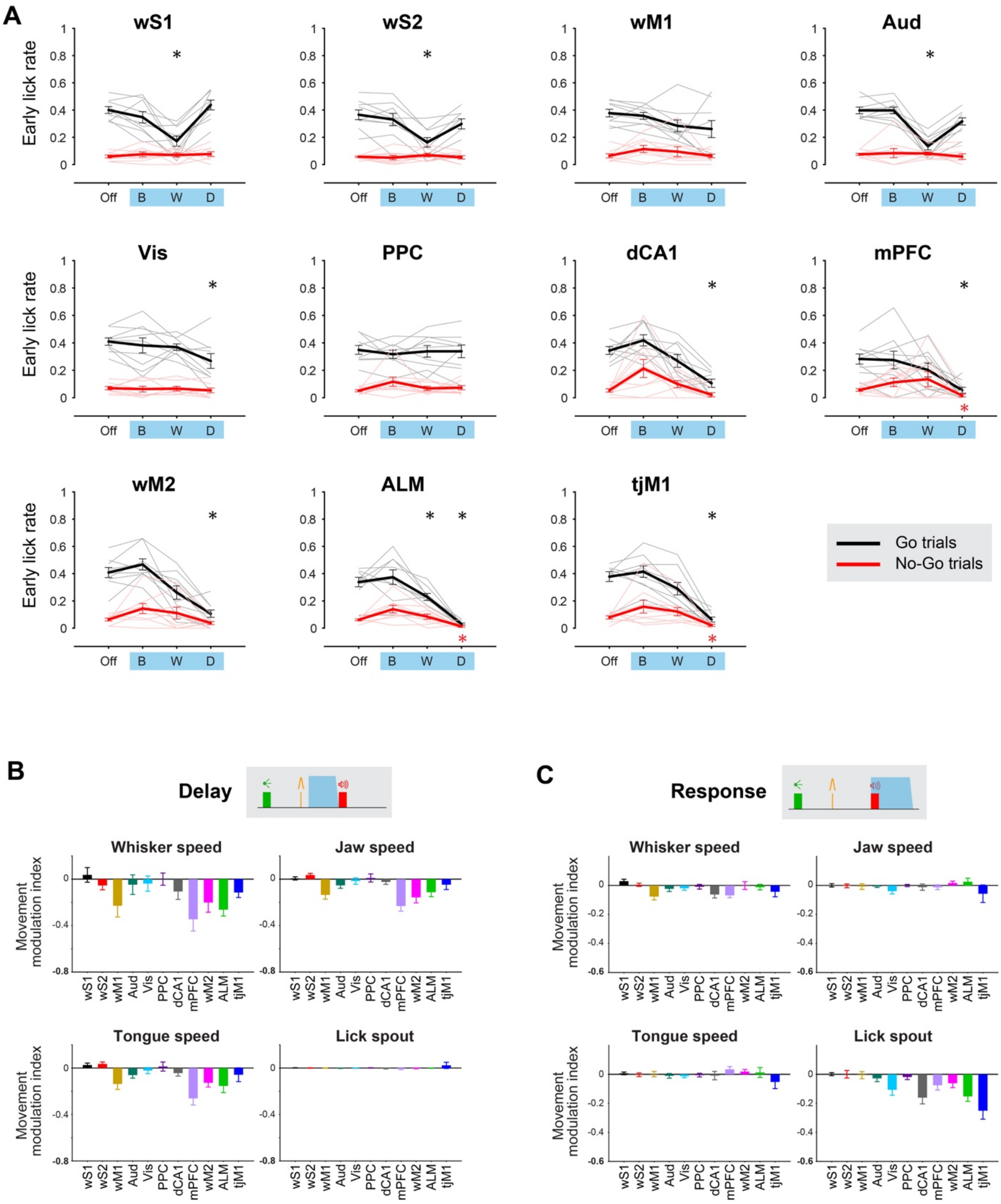

**Figure S8. Spatiotemporal effect of different regions on premature licking and movements, Related to Figure 7.**

(A) Behavioral impact of optogenetic inactivation across time windows for each brain region (mean ± SEM) on fraction of Early licks. For each area, Early lick rate in Go (black) and No-Go (red) trials are plotted for Light-off (Off), Baseline (B), Whisker (W), and Delay (D) windows. Asterisks represent significant difference comparing Early licks for light vs light-off trials in Go (black) or No-Go (red) trials (n=9 mice; *, $p<0.05$; Wilcoxon signed-rank test, Bonferroni correction for multiple comparisons).

(B) Changes in preparatory movements in the delay period. Change in orofacial movements (whisker, jaw and tongue speed) and licking pattern (lick spout sensor) during the delay period was quantified in trials with light (over the Delay) vs light-off trials (n=9 mice, mean ± SEM, see STAR methods for movement modulation index definition). For both light and no-light trials only Hit trials are included.

(C) Changes in movement during response window. Similar to (B), but when light was applied and movements were quantified during the response window. Similarly, only Hit trials are included for both light and light-off trials.

### Supplemental Table S1

| | | wS1 | wS2 | wM1 | Aud | Vis | PPC | dCA1 | mPFC | DLS | wM2 | ALM | tjM1 |
|---|---|---|---|---|---|---|---|---|---|---|---|---|---|
| Whisker | Novice | 0.56 | 0.56 | 0.32 | 0.06 | 0 | 0.05 | 0.12 | 0.08 | 0.14 | 0.17 | 0.15 | 0.06 |
| | Expert | 0.49 | 0.47 | 0.22 | 0.08 | 0.03 | 0.1 | 0.09 | 0.05 | 0.19 | 0.21 | 0.12 | 0.03 |
| | p | 0.332 | 0.106 | 0.029 | 0.681 | 0.13 | 0.192 | 0.579 | 0.294 | 0.392 | 0.223 | 0.288 | 0.0652 |
| Delay | Novice | 0 | 0.02 | 0.02 | 0.03 | 0 | 0.01 | 0 | 0.01 | 0.04 | 0.02 | 0.08 | 0.03 |
| | Expert | 0.06 | 0.08 | 0.07 | 0.04 | 0.02 | 0.05 | 0.02 | 0.04 | 0.1 | 0.12 | 0.24 | 0.08 |
| | p | 0.0558 | 0.0441 | 0.0394 | 0.702 | 0.287 | 0.119 | 0.441 | 0.0911 | 0.145 | 0.0000519 | 0.000000345 | 0.0203 |
| Lick initiation | Novice | 0 | 0.04 | 0.04 | 0 | 0.03 | 0.03 | 0 | 0.02 | 0.05 | 0.06 | 0.09 | 0.08 |
| | Expert | 0.04 | 0.04 | 0.04 | 0.02 | 0.06 | 0.02 | 0.04 | 0.02 | 0.08 | 0.08 | 0.11 | 0.09 |
| | p | 0.0996 | 0.932 | 0.71 | 0.21 | 0.378 | 0.774 | 0.218 | 0.958 | 0.52 | 0.24 | 0.273 | 0.628 |
| PostLick | Novice | 0 | 0.03 | 0.03 | 0.02 | 0.03 | 0.05 | 0 | 0.06 | 0.07 | 0.11 | 0.25 | 0.18 |
| | Expert | 0.05 | 0.07 | 0.07 | 0.03 | 0.05 | 0.01 | 0.07 | 0.03 | 0.1 | 0.12 | 0.18 | 0.2 |
| | p | 0.082 | 0.182 | 0.0936 | 0.537 | 0.509 | 0.0836 | 0.116 | 0.0369 | 0.566 | 0.753 | 0.0154 | 0.449 |
| LateLick | Novice | 0 | 0 | 0.02 | 0.06 | 0 | 0 | 0.03 | 0.02 | 0.07 | 0.02 | 0.17 | 0.07 |
| | Expert | 0.02 | 0.05 | 0.03 | 0.13 | 0.01 | 0.02 | 0.04 | 0.01 | 0.08 | 0.04 | 0.1 | 0.07 |
| | p | 0.277 | 0.0201 | 0.541 | 0.133 | 0.452 | 0.208 | 0.72 | 0.477 | 0.857 | 0.2 | 0.0138 | 0.872 |
| LED onset | Novice | 0.02 | 0 | 0.03 | 0.05 | 0.54 | 0.16 | 0.12 | 0.02 | 0.07 | 0.01 | 0.06 | 0.02 |
| | Expert | 0.05 | 0.02 | 0.02 | 0.03 | 0.6 | 0.34 | 0.05 | 0.03 | 0.04 | 0.02 | 0.02 | 0.01 |
| | p | 0.228 | 0.146 | 0.558 | 0.534 | 0.496 | 0.0042 | 0.175 | 0.427 | 0.302 | 0.739 | 0.000154 | 0.495 |
| LED delay | Novice | 0.02 | 0 | 0.03 | 0.03 | 0.19 | 0 | 0 | 0.01 | 0.02 | 0.01 | 0.04 | 0.01 |
| | Expert | 0.02 | 0 | 0.01 | 0.04 | 0.28 | 0.09 | 0.02 | 0.01 | 0.03 | 0.01 | 0.02 | 0 |
| | p | 0.889 | 1 | 0.102 | 0.856 | 0.17 | 0.0061 | 0.441 | 0.919 | 0.512 | 0.579 | 0.0418 | 0.0572 |
| Sound onset | Novice | 0.06 | 0.05 | 0.16 | 0.25 | 0.12 | 0.19 | 0.26 | 0.16 | 0.34 | 0.19 | 0.27 | 0.15 |
| | Expert | 0.16 | 0.18 | 0.16 | 0.6 | 0.06 | 0.29 | 0.13 | 0.12 | 0.41 | 0.14 | 0.2 | 0.14 |
| | p | 0.0472 | 0.00185 | 0.973 | 0.00000166 | 0.144 | 0.0947 | 0.0583 | 0.246 | 0.359 | 0.0732 | 0.0201 | 0.813 |
| Sound delay | Novice | 0.02 | 0.01 | 0 | 0.19 | 0.16 | 0.04 | 0.03 | 0 | 0.05 | 0.06 | 0.13 | 0.07 |
| | Expert | 0.05 | 0.06 | 0.06 | 0.19 | 0.07 | 0.07 | 0.07 | 0.04 | 0.15 | 0.04 | 0.14 | 0.05 |
| | p | 0.228 | 0.0555 | 0.00839 | 0.927 | 0.0609 | 0.322 | 0.393 | 0.0244 | 0.0501 | 0.329 | 0.793 | 0.425 |
| Jaw speed | Novice | 0.05 | 0.05 | 0.05 | 0.08 | 0.03 | 0.01 | 0.03 | 0.05 | 0.11 | 0.13 | 0.24 | 0.13 |
| | Expert | 0.1 | 0.13 | 0.17 | 0.13 | 0.17 | 0.12 | 0.1 | 0.08 | 0.25 | 0.17 | 0.24 | 0.26 |
| | p | 0.185 | 0.0299 | 0.000813 | 0.299 | 0.00518 | 0.00496 | 0.177 | 0.27 | 0.0219 | 0.18 | 0.891 | 0.0000915 |
| Tongue speed | Novice | 0.03 | 0.02 | 0.03 | 0.03 | 0.04 | 0.03 | 0.12 | 0.04 | 0.09 | 0.16 | 0.15 | 0.15 |
| | Expert | 0.05 | 0.1 | 0.09 | 0.04 | 0.07 | 0.06 | 0.07 | 0.07 | 0.12 | 0.11 | 0.15 | 0.23 |
| | p | 0.525 | 0.0158 | 0.0359 | 0.702 | 0.535 | 0.238 | 0.358 | 0.223 | 0.489 | 0.0701 | 0.788 | 0.0148 |
| Whisker speed | Novice | 0.6 | 0.38 | 0.45 | 0.28 | 0.4 | 0.28 | 0.5 | 0.35 | 0.79 | 0.61 | 0.34 | 0.31 |
| | Expert | 0.61 | 0.59 | 0.56 | 0.25 | 0.47 | 0.52 | 0.51 | 0.48 | 0.6 | 0.51 | 0.39 | 0.42 |
| | p | 0.962 | 0.000217 | 0.0296 | 0.662 | 0.326 | 0.000194 | 0.93 | 0.0089 | 0.0106 | 0.015 | 0.237 | 0.00421 |
| Previous Early Lick | Novice | 0.03 | 0.02 | 0.04 | 0.05 | 0.07 | 0.04 | 0.06 | 0.05 | 0.14 | 0.04 | 0.09 | 0.11 |
| | Expert | 0.06 | 0.06 | 0.09 | 0.01 | 0.05 | 0.04 | 0.02 | 0.03 | 0.03 | 0.08 | 0.06 | 0.07 |
| | p | 0.321 | 0.12 | 0.0684 | 0.105 | 0.5 | 0.95 | 0.186 | 0.448 | 0.000743 | 0.0272 | 0.0684 | 0.0706 |
| Previous Hit | Novice | 0.03 | 0.04 | 0.08 | 0.08 | 0.1 | 0.11 | 0.09 | 0.04 | 0.18 | 0.06 | 0.16 | 0.14 |
| | Expert | 0.09 | 0.12 | 0.12 | 0.02 | 0.04 | 0.05 | 0.08 | 0.07 | 0.06 | 0.08 | 0.09 | 0.1 |
| | p | 0.115 | 0.0239 | 0.287 | 0.0583 | 0.0955 | 0.0532 | 0.84 | 0.181 | 0.00456 | 0.291 | 0.0115 | 0.131 |
| Previous False-alarm | Novice | 0.06 | 0.04 | 0.09 | 0.09 | 0.13 | 0.11 | 0.12 | 0.04 | 0.16 | 0.08 | 0.18 | 0.16 |
| | Expert | 0.06 | 0.06 | 0.06 | 0.04 | 0.03 | 0.01 | 0.03 | 0.03 | 0.06 | 0.06 | 0.05 | 0.03 |
| | p | 0.923 | 0.548 | 0.257 | 0.0783 | 0.00963 | 0.000222 | 0.0257 | 0.503 | 0.00899 | 0.256 | 1.2E-09 | 4.69E-09 |
| Trial index | Novice | 0.41 | 0.48 | 0.6 | 0.52 | 0.53 | 0.52 | 0.68 | 0.63 | 0.52 | 0.54 | 0.57 | 0.59 |
| | Expert | 0.48 | 0.53 | 0.57 | 0.54 | 0.65 | 0.58 | 0.72 | 0.6 | 0.34 | 0.58 | 0.62 | 0.62 |
| | p | 0.369 | 0.389 | 0.519 | 0.718 | 0.0951 | 0.36 | 0.66 | 0.451 | 0.0168 | 0.366 | 0.191 | 0.472 |

**Supplemental Table S1. The fraction of neurons significantly modulated by different variables of the Poisson encoding model (GLM), Related to Figure 6.**

For the Poisson encoding model, the fractions of neurons in each brain region (*columns*) significantly modulated ($p<0.05$, likelihood ratio test) by different model predictors (*rows*) are shown separately for Novice (cyan) and Expert (purple) mice. Significant difference in the fraction of Novice vs Expert modulated neurons for each pair of brain region and model predictors are tested using the Pearson's chi-square test and the *p*-values are reported.
